# Using phage display for rational engineering of a higher affinity humanized 3’phosphohistidine-specific antibody

**DOI:** 10.1101/2024.11.04.621849

**Authors:** Gregory D. Martyn, Rajasree Kalagiri, Gianluca Veggiani, Robyn L. Stanfield, Indrani Choudhuri, Margaux Sala, Jill Meisenhelder, Chao Chen, Avik Biswas, Ronald M. Levy, Dmitry Lyumkis, Ian A. Wilson, Tony Hunter, Sachdev S. Sidhu

**Affiliations:** School of Pharmacy, University of Waterloo, 10 Victoria St A, Kitchener, ON, N2G 1C5, Canada; Molecular and Cell Biology Laboratory, Salk Institute for Biological Studies, La Jolla, CA, 92037, USA; Division of Biotechnology and Molecular Medicine, Louisiana State University, Baton Rouge, LA 70803, USA; Department of Pathobiological Sciences, School of Veterinary Medicine, Louisiana State University, Baton Rouge, LA, 70803, USA; Department of Integrative Structural and Computational Biology, Scripps Research, La Jolla, CA, 92037, USA; Laboratory of Genetics, Salk Institute for Biological Studies, La Jolla, CA, 92037, USA; Department of Physics, University of California San Diego, La Jolla, CA, 92093, USA; Center for Biophysics and Computational Biology, and Department of Chemistry, Temple University, Philadelphia, PA, 19122, USA; The Skaggs Institute for Chemical Biology, Scripps Research, La Jolla, California, 92037, USA

**Keywords:** Antibody Engineering, Protein Engineering, Phosphohistidine, Post-translational Modifications, Phage-Display

## Abstract

Histidine phosphorylation (pHis) is a non-canonical post-translational modification (PTM) that is historically understudied due to a lack of robust reagents that are required for its investigation, such as high affinity pHis-specific antibodies. Engineering pHis-specific antibodies is very challenging due to the labile nature of the phosphoramidate (P-N) bond and the stringent requirements for selective recognition of the two isoforms, 1-phosphohistidine (1-pHis) and 3-phosphohistidine (3-pHis). Here, we present a strategy for *in vitro* engineering of antibodies for detection of native 3-pHis targets. Specifically, we humanized the rabbit SC44-8 anti-3-pTza (a stable 3-pHis mimetic) mAb into a scaffold (herein referred to as hSC44) that was suitable for phage display. We then constructed six unique Fab phage-displayed libraries using the hSC44 scaffold and selected high affinity 3-pHis binders. Our selection strategy was carefully designed to enrich antibodies that bound 3-pHis with high affinity and had specificity for 3-pHis versus 3-pTza. hSC44.20N32F^L^, the best engineered antibody, has an ∼10-fold higher affinity for 3-pHis than the parental hSC44. Eleven new Fab structures, including the first reported antibody-pHis peptide structures were solved by X-ray crystallography. Structural and quantum mechanical calculations provided molecular insights into 3-pHis and 3-pTza discrimination by different hSC44 variants and their affinity increase obtained through *in vitro* engineering. Furthermore, we demonstrate the utility of these newly developed high-affinity 3-pHis-specific antibodies for recognition of pHis proteins in mammalian cells by immunoblotting and immunofluorescence staining. Overall, our work describes a general method for engineering PTM-specific antibodies and provides a set of novel antibodies for further investigations of the role of 3-pHis in cell biology.

**Significance Statement:** Histidine phosphorylation is an elusive PTM whose role in mammalian cell biology is largely unknown due to the lack of robust tools and methods for its analysis. Here we report the development of antibodies with unprecedented affinity and specificity towards 3-pHis and present the first crystal structures of a pHis peptide in complex with an antibody. Finally, we show how these antibodies can be used in standard molecular biology workflows to investigate pHis-dependent biology.

## Introduction

There are ∼600 known PTMs that underlie almost every process in cell biology (1). Protein phosphorylation is the best characterized PTM, with phosphorylation of serine (Ser; pSer), threonine (Thr; pThr) and tyrosine (Tyr; pTyr) being the most abundant and best characterized (2). Given the importance of PTMs in signal transduction pathways, it is unsurprising that enzymes involved in post-translational modification of proteins are frequently mutated in many diseases, such as cancer (3), and therefore represent attractive drug targets (4–6). Rare and non-canonical PTMs are emerging as additional key regulators of cell biology and biochemistry (7). Of particular interest is the role of phosphohistidine (pHis) in mammalian cell biology (8). This non-canonical PTM was first described in 1962 (9) and synthesized in 1966 (10), but despite a role for His phosphorylation being well characterized in prokaryotic cells (11), it remains understudied in mammalian cells. One of the major challenges in the detection and analysis of pHis is the acid and heat labile nature of the phosphoramidate (P-N) bond in pHis (12) and the lack of suitable reagents to study this PTM. Despite these obstacles, the discovery of histidine kinases and phosphatases (13), as well as modified experimental pipelines (14) and phosphoproteomics workflows (15), have recently enabled the study of this PTM in mammalian cells at a systems level scale.

Advances in the development of non-hydrolyzable pHis mimetics led to their subsequent use for raising anti-pHis antibodies (16–22). Some of the pivotal anti-pHis antibodies are rabbit-derived monoclonal antibodies (mAbs) that were obtained by immunizing with non-hydrolyzable phosphoryl-triazolylalanine (pTza; a pHis mimetic)-containing peptides (23). Subsequent structure-function analysis has provided insights into how these antibodies discriminate between 1- and 3-pTza in a peptide context (24). This suite of anti-pHis antibodies has been used by many groups to investigate the role of pHis in bacterial and mammalian physiology (25–29). Of these antibodies, the rSC44-8 3-pHis mAb has been most widely used as it has relatively higher affinity towards 3-pHis proteins. However, rSC44-8 is more specific for the 3-pTza analogue (K_D_: 0.24 nM) than the natural 3-pHis, where the rSC44-8 affinity is ∼75 fold lower (K_D_: 18 nM) (24).

The present study extends these advances by (i) humanizing the 3-pTza specific rabbit antibody, (ii) engineering the humanized antibody for improved affinity and specificity towards 3-pHis compared to 3-pTza, (iii) solving the first crystal structure of an antibody:3-pHis peptide complex, and (iv) demonstrating the use of the newly derived, high affinity anti-3-pHis antibodies to detect 3-pHis proteins in standard molecular and cell biology workflows. This advance was achieved by rational engineering of the human trastuzumab (4D5) antibody scaffold to accommodate rabbit complementarity determining regions (CDRs) by substituting specific variable region framework residues known as Vernier zone residues that modulate the conformation of CDRs (30, 31). We then used this humanized scaffold for constructing novel phage displayed antibody libraries and carried out affinity selections for both pTza- and pHis-containing peptides. Our engineering strategy allowed us to isolate new antibodies with increased affinity and specificity for 3-pTza- and 3-pHis-containing peptides. Furthermore, using novel, highly crystallizable Fab scaffolds (32–34), we solved 11 crystal structures of anti-3-pHis Fabs in complex with phosphorylated ligands, including the first antibody-pHis peptide structure. These structures in conjunction with quantum mechanical and molecular mechanical (QM/MM) studies provided insights into the recognition of 3-pTza versus 3-pHis by the humanized and engineered antibodies, explained the enhanced affinity of the engineered variant antibodies, and provide a basis for future engineering of antibodies towards specific pHis-peptides relevant to mammalian cell biology. Finally, we showed that the engineered 3-pHis specific antibodies identified biologically relevant pHis signals in mammalian cells using immunoblotting and immunofluorescence staining.

The significance of these studies is that they establish a generalizable strategy for engineering antibodies to recognize unstable non-canonical PTMs that can only be accomplished using *in vitro* evolution. Recent studies describing the engineering of a mouse monoclonal antibody for higher affinity recognition of pTyr highlight the importance of our approach (35). The enhanced affinity of anti-3-pHis antibodies can be used to elucidate the role of pHis in a variety of fields including cell biology, phosphoproteomics, signal transduction, and identifying novel drug targets (36).

## Results

### Rabbit Antibody Scaffold Engineering and Humanization

To engineer 3-pTza specific rabbit antibodies for improved affinity and specificity for 3-pHis, we elected to use SC44-8 (hereafter referred to as rSC44), which is a 3-pHis specific rabbit mAb that has partial sequence selectivity for Gly-3-pHis-Ala sites and binds to an AGAG-3-pHis-AGAG peptide with ∼20 nM affinity. We first assessed whether rSC44 could be produced as IgG and Fab proteins from mammalian cells and displayed as Fab and scFv versions on M13 bacteriophage. The rSC44 antibody could reliably be produced as IgG and Fab proteins, as previously described (24), but could not be displayed on phage in either Fab or scFv formats. This led to an extensive rational engineering campaign on the rSC44 scaffold to humanize it for directed evolution campaigns using phage display.

We first sought to fully humanize the rabbit antibody into the human 4D5 framework (trastuzumab), which displays efficiently on phage (37). Humanization of rabbit antibodies has traditionally been accomplished using CDR grafting and substitution of specific Vernier zone residues in the variable framework (38, 39). The sequences of rSC44 were aligned with those of the human 4D5 scaffold using Kabat (40) and IMGT nomenclatures (41) for both the variable light (V_L_) and variable heavy (V_H_) regions (Fig. S1), prior to loop grafting of all 6 rSC44 CDRs into the human 4D5 scaffold using both IMGT and Kabat boundaries. These humanized versions displayed on phage but did not bind the 3-pTza antigen. We then identified sets of Vernier zone residues that could be important for CDR orientation and therefore antigen binding. Several successive rounds of combinatorial back mutation were necessary to engineer a functional human scaffold for rSC44. Hereafter, we will refer to specific amino acid positions using Kabat nomenclature (Fig. S1*C,D*).

The humanized version of rSC44 (hSC44) had 2 and 14 Vernier zone residues in the V_L_ and V_H_ domains back mutated to rabbit sequences, respectively (Fig. 1*A*) and could be produced as IgG and Fab proteins (Fig. S1*E*). Furthermore, the IgG version of hSC44 bound the 3-pTza peptide with high affinity (K_D_ = 1.2 nM) but exhibited no binding to the 1-pTza or the non-phosphorylated (No PO_4_ ^-^) version of the peptide as measured by biolayer interferometry (BLI; Fig. 1*B*). Finally, Fab and scFV versions of hSC44 displayed on phage could bind 3-pTza as measured by ELISA (Fig. S1*F*).

**Fig. 1.**
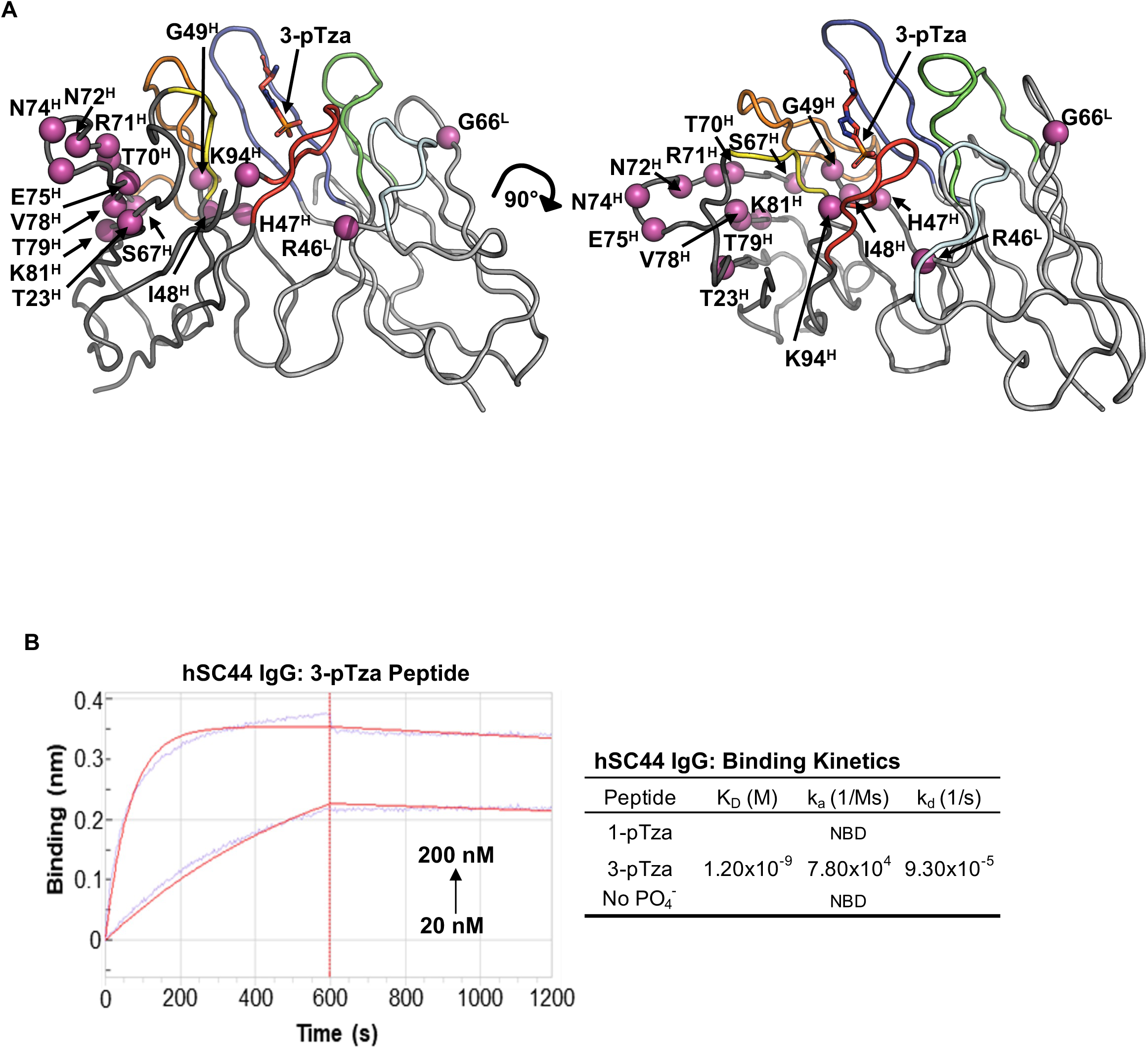
Humanization of the rabbit (rSC44) antibody. *(A)* Structure of rSC44 (PDB ID: 6X1V) in complex with a 3-pTza containing peptide (3-pTza shown as sticks colored in red) was used to guide humanization. The V_H_ and V_L_ domains are colored dark and light grey respectively. The CDRs are colored as follows: H3, red; H2, orange; H1, yellow; L3, blue; L2, cyan; L1, green. Residues in important framework positions, including known Vernier Zone residues are colored magenta and represented as spheres highlighting their Cα position. Residue numbering is in accordance with Kabat nomenclature. *(B)* Biolayer Interferometry (BLI) data for hSC44 IgG binding to 3-pTza, 1-pTza and a non-phosphorylated (no PO ^-^) peptide. The table shows the binding constants determined from BLI traces, and sample traces are shown for hSC44 binding to the immobilized 3-pTza peptide. NBD indicates no binding detected.

### Phosphohistidine Peptide Immunoprecipitation

To generate 1/3-pHis-containing peptides for antibody selection, we developed a custom immunoprecipitation workflow (Fig. S2). Biotinylated-His-peptides were incubated with potassium phosphoramidate for 16 h in 1x HEPES buffered saline (HBS; pH = 8.5) to generate pHis peptides. We then constructed a rProteinA-IgG resin using the rSC1 (1-pTza specific) and rSC44 (3-pTza specific) IgG’s described previously (23), in order to enrich either 1-pHis or 3-pHis peptides, respectively. Elution conditions using 2 M imidazole (pH 8.5) were optimized and eluates were analyzed using ELISA (see Materials and Methods for a detailed description).

### Phage Displayed Library Construction and Affinity Selections

In an attempt to both increase hSC44 affinity and improve specificity for 3-pHis versus 3-pTza peptides, we designed and constructed 6 unique phage displayed Fab libraries (Fig. S3-4). To aid library design, we analyzed the previously reported structure of rSC44 in complex with a 3-pTza peptide (PDB ID: 6X1V) and selected for diversification residues in both V_L_ and V_H_ regions. Libraries 1-5 (hSC44 1-5) diversified residues that are in direct contact with the 3-pTza side chain or have side chains oriented towards and proximal (within 10 Å) to the 3-pTza side chain (24). Library 6 diversified key Vernier zone and additional framework residues that were involved in CDR coordination and were important for creating a stable human scaffold. All libraries contained 10^9^-10^10^ unique variants and were generated using a “soft randomization” strategy that favors wild-type (wt) sequence but allows for diversity across all positions sampled (Figs. S3 and S4, Table S1) (42).

To select for Fab variants with high affinity and specificity for 3-pHis peptides, but not the 1-pHis isoform, we devised a selection strategy in which 1-pTza and non-phosphorylated peptides (No PO_4_ ^-^) were used to deplete all 6 libraries of cross-reactive Fabs (pools a-d, Fig. 2*A*, Table S2). For example, pools a and b were positively selected for 3-pTza whereas pools c and d were positively selected for 3-pHis. Pools b and d were subjected to negative selections against 1-pTza. Selections were carried out with pre-clearing steps on BSA and streptavidin as well as stringent negative selections on non-phosphorylated peptides. All 6 hSC44 libraries were cycled through 5 rounds of affinity selections and yielded 32 unique sequences (Fig. 2*B*). No hSC44 template was detected across pools a-d, indicating the successful progression of affinity selections. We produced each unique variant as a phage displayed Fab as described previously (43), normalized their concentrations, and assessed binding to 1/3-pTza and 1/3-pHis peptides relative to hSC44 using high-throughput ELISAs. Thirty of the 32 unique sequences displayed increased binding to either 3-pTza or 3-pHis and no binding to 1-pTza or 1-pHis (Fig. 2*B*). Remarkably, we achieved high 3-pHis specificity in our selection campaigns, as all the enriched antibodies that were positively selected for 3-pHis showed preferable binding to 3-pHis over the phospho-mimetic 3-pTza embedded in the same peptide sequence.

**Fig. 2.**
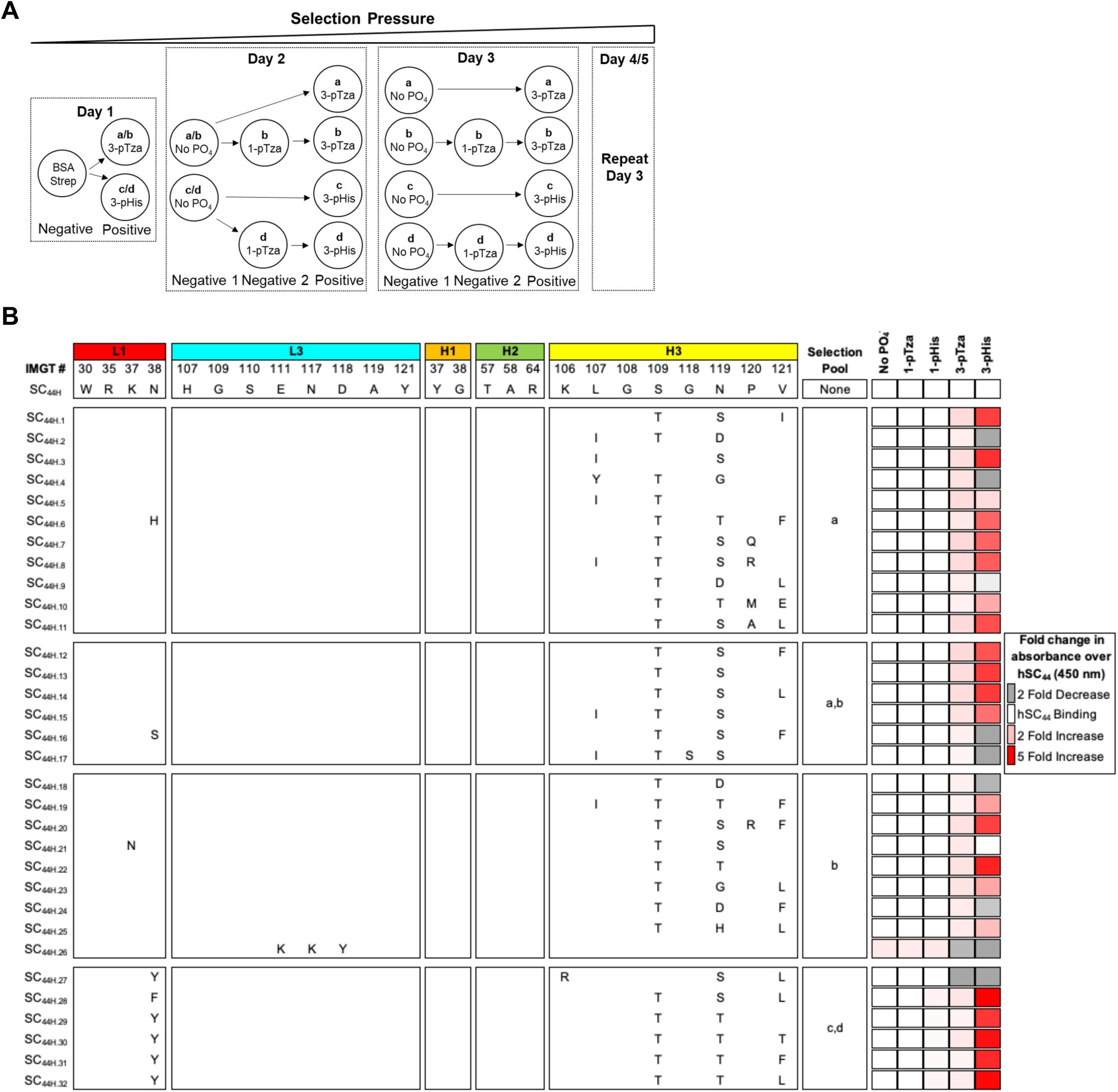
Outline of affinity selections protocol and selection of 3-pHis-specific hSC44 variants. *(A)* Schema depicting affinity selection strategy. Libraries were combined on day 1 with unique outputs from different positive selections carried through the remainder of the selection process. The strategy resulted in 4 unique pools labelled a-d which represent unique positive selections. No PO_4_ is a non-phosphorylated version of the His-peptide. *(B)* hSC44 variants isolated after 5 rounds of affinity selections. The amino acid sequence (Kabat nomenclature) of hSC44 residues randomized in different libraries are shown at the top. Sequences in hSC44 variants that were the same as hSC44 are blank and differences are shown with the single-letter amino acid code. The pool, which each unique variant was identified in, is listed next to each sequence. Unique variants were reamplified as Fab-Phage, normalized for concentration and assayed by phage-ELISA for binding to non-phosphorylated (No PO_4_^-^), 1-pTza, 1-pHis, 3-pTza and 3-pHis peptides. Binding is shown as fold change in absorbance over hSC44.

Sequence analysis of the selected 3-pTza and 3-pHis antibodies revealed that most of the substitutions resided in CDR-H3, with almost all variants displaying S97T^H^ and N99T/S^H^ substitutions (Fig. 2*B*). These substitutions were correlated with increased binding to 3-pTza and/or 3-pHis. Furthermore, we observed that while N99^H^ was substituted with D, H or G in the CDR-H3 of Fabs selected for 3-pTza-containing peptides, these substitutions were absent in Fabs positively selected for 3-pHis and correlated with a moderately increased affinity for 3-pTza and diminished or no binding to 3-pHis. Some variants had a V100aL/F^H^ substitution in CDR-H3, but this substitution was less frequent and did not correlate with enhanced binding. Finally, the N32Y/F^L^ substitution in CDR-L1 was observed only in variants selected for 3-pHis and conferred preferential binding to 3-pHis over 3-pTza (Fig. 2B right hand panels). Together, these results indicate that a small set of substitutions in CDR-H3 can increase affinity for both 3-pTza and 3-pHis, and that a single substitution in CDR-L1 contributes to enhanced binding of 3-pHis over 3-pTza.

### hSC44 Variant Affinity, Specificity, and Developability

To investigate the antibody variants in greater detail we expressed them as IgGs (Fig. S5*A,B*). Twenty-eight of the 32 variants had higher yields than hSC44, 17 of which were even greater than the high-yield trastuzumab benchmark. We selected 7 hSC44 variants that bound 3-pTza/3-pHis with high apparent affinity and had favorable IgG yields and measured their EC_50_ values for the 3-pHis peptide to estimate their affinities using ELISA (Fig. S5*C*). In agreement with the phage ELISA screening (Fig. 2*B*), all variants had EC_50_ values that were up to 2-fold lower than hSC44. We observed that all isolated antibodies displayed a substantial improvement in binding to 3-pHis relative to the parental hSC44, with hSC44.20 showing the highest apparent affinity (Fig. S5*C*).

We further investigated the affinity of 8 antibody variants with the greatest increase in apparent affinity for 3-pTza/3-pHis using biolayer interferometry (BLI, Table 1 and Fig. S6). BLI measurements demonstrated that hSC44 variants positively selected for the 3-pTza peptide displayed affinities in the 100-1000 pM range, with one variant (hSC44.20) reaching the detection limit of the instrument (<1x10^-12^ M) compared with 1.2 nM for parental hSC44. Variants that were positively selected for 3-pHis (hSC44.28 and hSC44.31) had a small increase or decrease in affinity for 3-pTza compared with hSC44, respectively.

**Table 1.** Affinity measurements of hSC44 variants against 3-pTza peptides using bio-layer interferometry.

| Name | 3-pTza Peptide |  |  |
| --- | --- | --- | --- |
| | $K_D$ (M) | $k_a$ (1/Ms) | $k_d$ (1/s) |
| hSC44.1 | 5.88E-10 | 1.41E+05 | 8.32E-05 |
| hSC44.3 | 3.52E-10 | 4.07E+04 | 1.43E-05 |
| hSC44.8 | 1.85E-10 | 3.59E+05 | 6.66E-05 |
| hSC44.13 | 4.05E-10 | 1.82E+05 | 7.37E-05 |
| hSC44.20 | <1.0E-12 | 2.57E+05 | <1.0E-07 |
| hSC44.20.N32F<sup>L</sup> | 1.54E-09 | 3.73E+04 | 5.73E-05 |
| hSC44.20.N32Y<sup>L</sup> | <1.0E-12 | 3.51E+04 | <1.0E-07 |
| hSC44.22 | 4.95E-10 | 9.06E+04 | 4.49E-05 |
| hSC44.28 | 1.21E-09 | 2.49E+04 | 3.00E-05 |
| hSC44.31 | 2.71E-09 | 1.47E+04 | 3.98E-05 |

Given that hSC44.20 had the highest affinity for 3-pTza and that all antibody variants isolated using 3-pHis as bait possessed a N32Y/F^L^ in their CDR-L1, we decided to shuffle the light chain of hSC44.20 to obtain hSC44.20.N32Y^L^ and hSC44.20.N32F^L^. Affinity measurements for 3-pTza revealed that, while the N32Y^L^ substitution did not affect binding to 3-pTza (K_D_ <1x10^-12^ M), the N32F^L^ substitution substantially reduced 3-pTza affinity (K_D_ <1.54x10^-9^ M, Table 1, Fig. S6). We then measured the binding affinity of hSC44.20.N32Y^L^ and hSC44.20.N32F^L^ for 3-pHis and showed that both variants bound 3-pHis peptides with affinities higher (N32F^L^ = 1.6 nM and N32Y^L^ = 9.5 nM) than the parental antibody (Table 2, and Fig S7). No binding to pTyr, pSer, pThr or the non-phosphorylated version of the peptide could be detected for any of the variants by ELISA (Fig. S5*D*).

**Table 2.** Affinity measurements of hSC44 variants against 3-pHis peptides using bio-layer interferometry.

| Name | 3-pHis Peptide |  |  |
| --- | --- | --- | --- |
| | $K_D$ (M) | $k_a$ (1/Ms) | $k_d$ (1/s) |
| hSC44.20 | 2.10E-08 | 7.55E+04 | 1.60E-03 |
| hSC44.20.N32F<sup>L</sup> | 1.68E-09 | 1.39E+04 | 1.64E-05 |
| hSC44.20.N32Y<sup>L</sup> | 9.50E-09 | 8.88E+03 | 2.61E-05 |

To assess the developability of the new antibody variants, we performed a specificity ELISA against known antigenic molecules, as cross-reactivity is indicative of poor developability (44) (Fig. S8). We did not detect binding to any of the molecules tested, indicating no cross-reactivity. Together these, results demonstrate that humanized hSC44 and its engineered variants provide a scaffold that can produce IgGs with high affinity for 3-pHis and are correlated with good developability for future engineering and use in research pipelines.

### Antibody Structure Determination

To better understand the structure-function relationships of antibody:pHis binding, we undertook an extensive crystallization campaign to solve the structures of Fab versions of hSC44, hSC44.20, and hSC44.20.N32F^L^. Initial crystallization attempts with hSC44 Fab failed to produce any crystals. To overcome this issue, we utilized a method that enhances the crystallization of Fabs by substituting several residues distal to the antibody paratope that favor crystal packing interfaces. This method was based on (i) altering the constant kappa (cκ) light chain β-sheet (34) along with two substitutions upstream of this site (32), which are collectively known as “S1C” (Fig. S9), and (ii) substituting the elbow region residues of the heavy chain (33) (Fig. S10; Table S3). The S1C substitutions were introduced either alone or in combination with the elbow substitutions (collectively called “S1CE” substitutions) into the scaffolds described above and used in crystallization trials to solve structures in the apo, 3-pTza or 3-pHis complexed forms. These substitutions enabled us to obtain dozens of new crystals and to solve 11 new structures, including three 3-pHis peptide/Fab co-crystal structures (Tables S4-5). Notably, the substitutions that enhanced crystallization did not affect the overall structure of the antibody or the epitope.

### Effect of Vernier Zone Residue Back Mutation on hSC44 Structure

Humanization of the rSC44 antibody required back mutation of 2 Vernier zone residues in the human light chain and 14 residues (Vernier plus additional framework residues) in the heavy chain (Fig 1*A*; Fig. S1). To evaluate the structural similarity of rSC44 and hSC44, we performed an alignment of the Cɑ coordinates of the V_H_ region, which showed a root mean square deviation (RMSD) of 0.5 Å (Fig. S11*A*). Most of the Vernier zone residue changes are structurally justified as they are involved in positioning the CDR loops. For example, the L46R^L^ back mutation in the light chain is important, as R46^L^ interacts with the main chain of N99^H^ and P100^H^ on the CDR-H3 loop that is relevant for coordinating the phosphoryl group in 3-pTza (Fig. S11*B*). Most of the substituted Vernier residues in the heavy chain are located in framework regions 2 and 3 (Fig. *1A*; Fig. S1). Residue F34^H^ is involved in forming hydrophobic interactions with F27^H^, L4^H^, K94^H^ and I102^H^. While these residues form the core of the antibody, K94^H^ is involved in hydrogen bonding with the phosphoryl group (Fig. S11*C*). Substitutions from ^47^WVAS^50^ to ^47^HIGY^50^ in the framework region 2 and start of CDR-H2 of the heavy region are critical, as this loop is at the interface between the heavy chain and light chain variable domains. While H47^H^ establishes hydrogen bond interactions, Y50^H^ forms T-shaped pi-pi stacking with Y96^L^ (distance 5.0 Å, angle 75°, and offset 1.5 Å) at the interface, which in turn interacts with the phosphoryl moiety (Fig. S11*D*). Residues ^61^SWAK^64^ in CDR-H2 of the heavy chain are directed towards the core of the antibody and structurally support the ^47^HIGY^50^ strand (Fig. S11*D*). Residues ^70^TRNTNENTV^78^ on framework region 3 of the heavy chain support the orientation of the CDR-H1 and CDR-H2 loops. Together, the Vernier zone substitutions uphold the structure of the paratope that interacts with either 3-pHis or 3-pTza containing epitopes.

### Structural Analysis of 3-pHis Recognition by hSC44, hSC44.20, and hSC44.20.N32F^L^

In order to understand the structural basis for recognition of 3-pHis versus 3-pTza by hSC44 variants, we co-crystallized them with 3-pTza and 3-pHis peptides. Though 3-pHis peptides are labile and undergo spontaneous hydrolysis, our previous study indicated that bound rSC44-8 shields the 3-pHis moiety and slows down its hydrolysis (24). Inclusion of the entire phosphorylation reaction together with the hSC44 variant proteins in the co-crystallization mix allowed the antibodies to bind to and protect 3-pHis peptides during crystallization leading to crystals containing intact 3-pHis peptides. We aligned co-crystal structures of hSC44, hSC44.20 and hSC44.20.N32F^L^ in complex with a 3-pTza or 3-pHis peptide using the Cɑ coordinates of the V_H_ domain and demonstrated that these structures are highly similar to one another with RMSD’s <1.0 Å (Fig. S12). Consequently, the peptide location in the CDR regions of the hSC44 variants are nearly identical to one another as well as in rSC44 (Fig. 3*A-C* and S11*A*). The phosphoryl moiety of 3-pHis forms a hydrogen bond network with K94^H^ and the CDR-H3 residues 95-99 (LGSGN^H^ in hSC44, LGTGS^H^ in hSC44.20 and hSC44.20.N32F^L^) that form the canonical phosphoryl-recognizing GXGX motif (45) (Fig. 3*D-F*). The GXGX motif forms a loop around the phosphoryl group in the shape of a ‘nest’ and interacts through the main chain amides of G96^H^ and G98^H^, and the side-chain hydroxyl of S/T97^H^ and side-chain or backbone interactions with N/S99^H^. While N32^L^ in CDR-L1 of hSC44.20 makes water mediated interactions with the phosphoryl group, the same interaction is absent in hSC44.20.N32F^L^ due to the substituted Phe at this position (Fig. 3*F*). Additionally, the Y96^L^ hydroxyl group from CDR-L3 in all variants also makes direct hydrogen bonds with the phosphoryl group. This interaction is further strengthened by water-mediated hydrogen bonds to G33^H^ and Y50^H^.

**Fig. 3.**
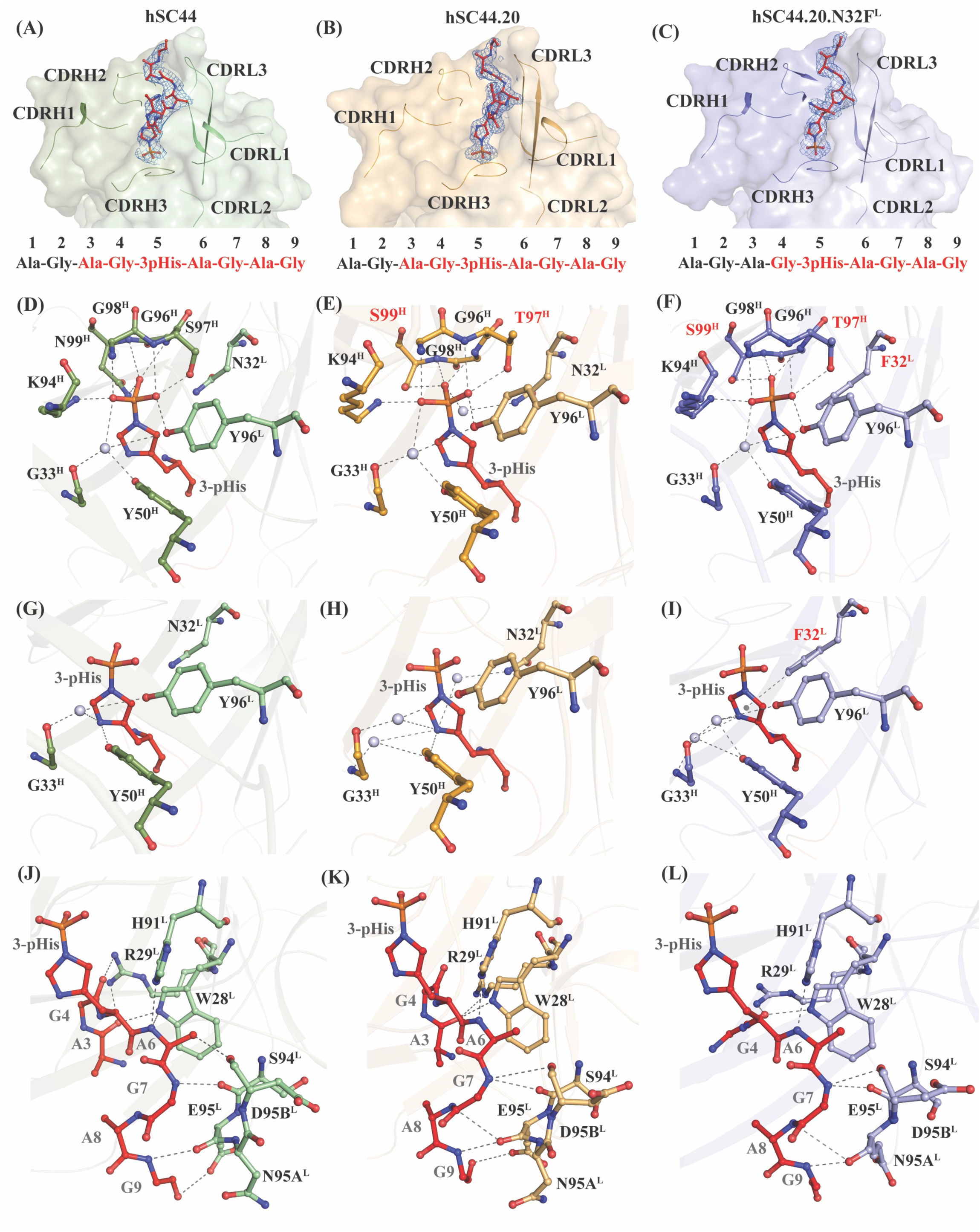
Structural analysis of hSC44 variants bound to 3-pHis. *(A-C)* Omit maps (*F_O_-F_C_* map) of *(A)* hSC44 Fab (green), *(B)* hSC44.20 Fab (orange), and *(C)* hSC44.20.N32F^L^ Fab (blue) bound to a 3-pHis peptide (red). The peptide is contoured at 1.0 σ and the CDR loops are represented as cartoon under the surface. The sequence of the peptide used for crystallization is shown below and the residues in red font have interpretable electron density. *(D-F)* Hydrogen bond network of phosphoryl moiety contacts with CDR residues and coordinating water molecules (grey spheres) of *(D)* hSC44, *(E)* hSC44.20, and *(F)* hSC44.20.N32F^L^. Substituted residues in *(E-F)* that are different from hSC44 (*D)* are shown in red. *(G-I)* Hydrogen bond network of imidazole ring contacting residues and coordinating water molecules of *(G)* hSC44, *(H)* hSC44.20, and *(I)* hSC44.20.N32F^L^. *(J-L)* Hydrogen bond network of CDR residues contacting the ligand backbone and non-3-pHis side chain atoms in *(J)* hSC44, *(K)* hSC44.20, and *(L)* hSC44.20.N32F^L^.

In 3-pHis, the N3 nitrogen is phosphorylated, enabling the N1 nitrogen in the imidazole ring to interact with specific residues lining the binding pocket. The N1 nitrogen forms either direct or water-mediated hydrogen bond interactions with G33^H^, Y50^H^ and Y96^L^ in all hSC44 variants (Fig. 3*G-I*). The substitution N32F^L^ in hSC44.20.N32F^L^ increased the affinity of the antibody for the 3-pHis peptide by ∼10-fold. This is likely due to the formation of T-shaped pi-pi stacking interactions with the imidazole side chain that are lacking in other variants (Fig. 3*I*). We further investigated the effect of N32F^L^ substitution using Quantum Mechanics/Molecular Mechanics (QM/MM)-based contact analysis which revealed that though hSC44.20 and hSC44.20.N32F^L^ have equal numbers of contacts with the 3-pHis peptide, the N32F^L^ substitution introduced a new hydrophobic interaction (Fig. S13). Inspection of the structure following local alignment reveals a 1.2 Å shift of the imidazole ring closer to the N32F^L^ substitution when 3-pHis is bound to hSC44.20.N32F^L^, relative to the position of the peptide when it is bound to hSC44.20 (Fig. S14*A*). A more detailed QM/MM analysis indicates that the side chain of the N32F^L^ substitution establishes a T-shaped pi-pi interaction with the imidazole ring in pHis (Fig. S14*B*), which likely stabilizes the observed shift in peptide position when bound to hSC44.20.N32F^L^. The resulting stable π-π interaction explains the increased binding affinity of hSC44.20.N32F^L^ to 3-pHis (Fig. S15 and Table S6).

Ordered electron density was found for the AG-3-pHis-AGAG peptide in the variants of hSC44, in keeping with the pseudo-sequence specificity or partial-sequence dependence property of rSC44 (24). CDR-L1 W28^L^ and R29^L^ interact with the N-terminal residues of the peptide (A3 and G4) while CDR-L3 residues (H91^L^, S94^L^, E95^L^, N95a^L^ and D95b^L^) make hydrogen bond interactions with the peptide backbone at its C-terminus (G7-G9) (Fig. 3*J-L*). This network of hydrogen bonds is retained among hSC44 variants and is implicated in the higher affinity and specificity for the peptide ligand. This is further corroborated by thermal shift assays (Fig. S16A) and BLI assays (Fig. S16B) where the hSC44 variants show specificity for 3-pHis or 3-pTza-containing peptides of ACLYana (AGAG-pHis-AGAG), ACLY (VQFG-pHis-AGAC) and SCS (RRMG-pHis-AGAI), but do not bind other 3-pHis or 3-pTza peptides (Fig. S16).

### In-Depth Structure-Function Analysis of hSC44.20.N32F^L^ Substitutions

The high affinity variant hSC44.20.N32F^L^ has four substitutions in CDR-H3 (S97T^H^, N99S^H^, P100R^H^ and V100aF^H^) and one in CDR-L1 (N32F^L^) that differentiate it from the parental hSC44. These residues do not affect the conformations of their respective CDR-H3 loops and are essential for phosphoryl coordination of the pHis moiety, with T97^H^ and S99^H^ making hydrogen bonds with the phosphoryl group (Fig. 4 middle-inset). These residues also stabilize the CDR-H3 loop by forming intra- and inter-molecular networks with other CDRs as well as framework residues (Fig. 4*A,C*). S99^H^ forms hydrogen bonds with K94^H^ and R46^L^ (Fig. 4A). The guanidino group of R100^H^ in hSC44.20 forms hydrogen bond interactions with Y49^L^ and S56^L^, which otherwise could not be formed with P100^H^ of hSC44 (Fig. 4B). The methyl group on T97^H^ in hSC44.20 is directed towards the core of the antibody and makes a hydrophobic interaction with V89^L^ (distance 4.2 Å) and Y96^L^ (distance 3.5 Å) (Fig. 4*C*). F100a^H^ forms a T-type pi-pi stacking interaction with Y32^H^ (distance 5.5 Å, angle 68.0°) and F27^H^ (distance 4.4 Å, angle 86°) (Fig. 4D). The hSC44.20.N32F^L^ variant is selective for 3-pHis while maintaining high affinity (Tables 1-2). This variant has an additional N32F^L^ substitution in CDR-L1 that was specifically selected for in 3-pHis selections (Fig. 2*B*). This additional substitution forms T-shaped pi-pi stacking interactions with W28^L^ (distance 5.9 Å, angle 81°) and the imidazole ring (distance 4.2 Å, angle 86°) of the 3-pHis peptide. These pi-pi interactions mediated by F32^L^ may help to stabilize the CDR-L1 loop, resulting in selectivity for 3-pHis ligand (Fig. 4*E*).

**Fig. 4.**
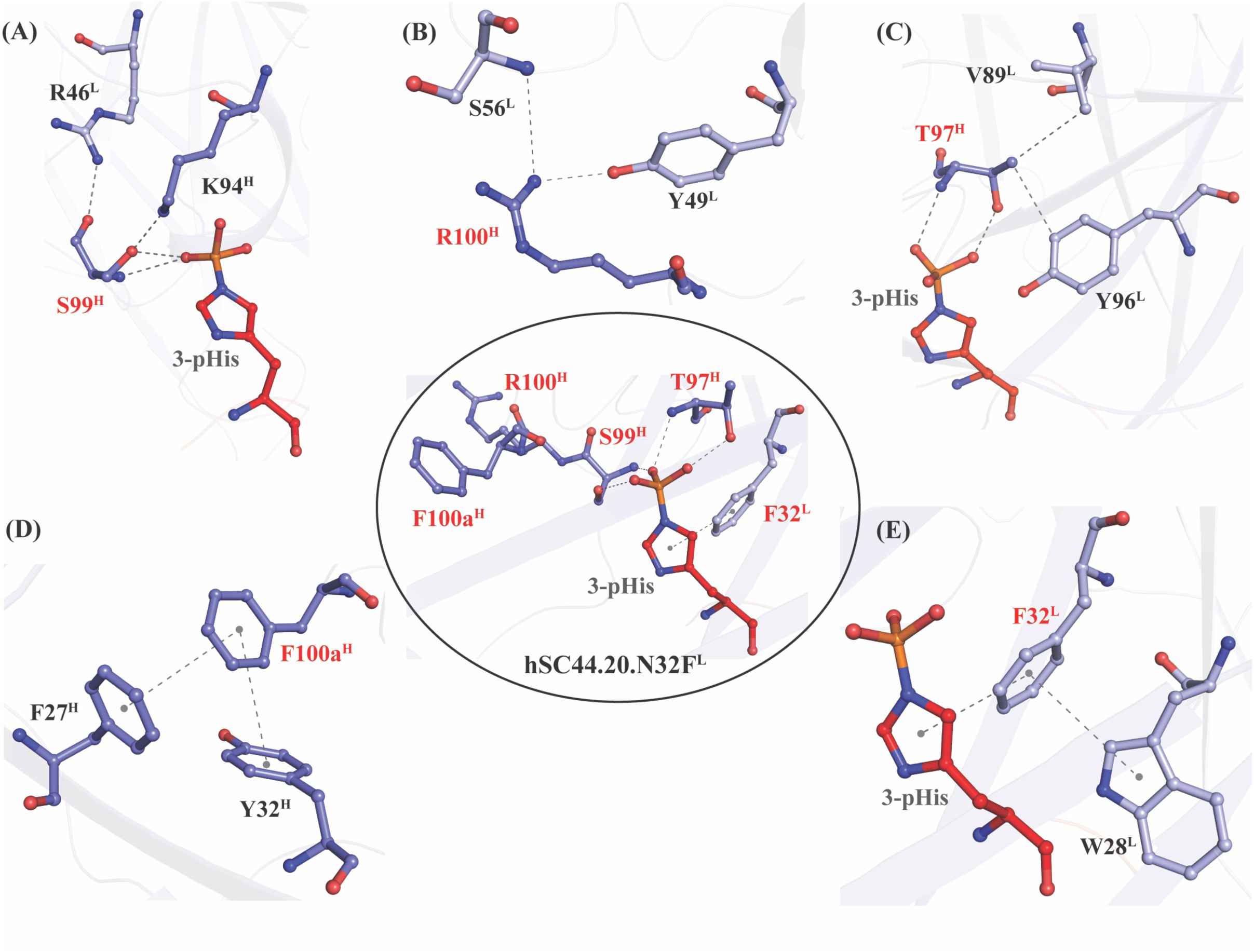
Structural analysis of effects of hSC44.20.N32F^L^ substitutions relative to hSC44 on 3-pHis recognition. The middle-inset shows the structure of hSC44.20.N32F^L^ Fab and the 3-pHis side chain of the peptide ligand, and it highlights the CDR-H3 and CDR-L1 residues (red font) that differ from hSC44. *(A)* The sidechain of S99^H^ makes hydrogen bonds with the phosphoryl group of 3-pHis. The sidechain and backbone residues interact with proximal framework residues R46^L^ and K94^H^. *(B)* The sidechain of R100^H^ points away from 3-pHis making contacts with framework residues S56^L^ and Y49^L^. *(C)* The sidechain and backbone atoms of T97^H^ make contact with the phosphoryl moiety of 3-pHis. The methyl group of the T97^H^ sidechain makes hydrophobic contacts with framework residues V89^L^ and Y96^L^. *(D)* The F100a^H^ side chain makes a sandwiching T-shaped π-π interaction with the side chains of framework residues F27^H^ and Y32^H^. *(E)* The F32^L^ side chain also makes a sandwiching T-shaped π-π interaction with the imidazole ring of 3-pHis as well as the side chain of CDR-L1 residue W28^L^.

### Analysis of Differences between 3-pTza and 3-pHis Coordination

Although hSC44 antibody variants are structurally similar, their affinity towards the 3-pTza peptide is >1000 times more than 3-pHis peptide (Table 1 and 2). To understand these differences, we analyzed the structures of hSC44 variants bound to 3-pTza or 3-pHis in greater detail (Fig. S17). This analysis highlights the high structural similarity of hSC44 variants to one another (RMSD <1 Å between Cɑ atoms in the V_H_ domain) when bound to either 3-pTza or 3-pHis. For simplicity, we further analyzed the hSC44.20 variant in complex with 3-pHis versus 3-pTza peptides (Fig. S18*A,B*). The CDR residues that interact with either 3-pTza or 3-pHis are similar in both structures (Fig. S18*C-F*). A water molecule buried at the interface between the peptide and CDR of the Fab, which mediate antibody-antigen interactions, is conserved between the 3-pTza and 3-pHis complexes (Fig. S18*C,D*). The anticipated difference between the 3-pTza and 3-pHis complexes mainly involves the triazolyl vs imidazole ring interactions. The triazolyl ring in the 3-pTza has three nitrogen groups (at ring positions 1, 2 and 5, with a phosphoryl group attached to the carbon at position 3) whereas 3-pHis has two nitrogens (at ring positions 1 and 3, with a phosphoryl group linked to the nitrogen at position 3 (Fig. S19*A*). The N1 atom of the 3-pHis sidechain ring and the N1 atom in the corresponding position of the pTza triazolyl ring both form water-mediated interactions with G33^H^ and Y50^H^ (Fig. S19*B,C*). The nitrogen atom at the triazolyl N2 position forms hydrogen bonds to K94^H^ and S99^H^ that are absent in the 3-pHis peptide, as it contains a carbon atom at this corresponding position (Fig. S19*B*). However, the >1000 fold difference in affinity observed for 3-pTza versus 3-pHis bound to hSC44 variants could not be explained by crystal structures. Hence, we carried out QM/MM calculations to rationalize this affinity difference between native and mimetic peptide. Molecular mechanics (MM)-based contact analysis reveals that, in comparison to 3-pHis, the 3-pTza moiety forms a greater number of contacts with the paratope of hSC44.20, and the stable contacts are predominantly associated with the phosphoryl moiety (Fig. S20). The chemical properties around the phosphoryl moiety differ between the two peptides. In 3-pHis, the phosphoryl moiety is attached via a nitrogen atom, whereas in 3-pTza, the phosphoryl moiety is attached via a carbon atom (Fig. S21*A*). Nitrogen, being more electronegative than carbon (46) (electronegativity 3.04 versus 2.55 for Nitrogen or Carbon, respectively), pulls more electron density from the phosphoryl moiety in 3-pHis (Fig. S21*B-D*), weakening its contacts with the surrounding environment. This accordingly leads to reduced binding affinity of 3-pHis to hSC44.20 in comparison to 3-pTza.

### Apo Structures of hSC44 Variants

The CDRs of the hSC44 variants are highly positively charged (24) and as a result, crystallization trials with hSC44 in the peptide-free state yielded structures with different small anionic groups bound in the paratope that were acquired during the purification process or from the crystallization buffers. These moieties include a citrate ion in hSC44, a phosphate ion or glutamic acid (from a symmetry-related Fab) in hSC44.20, and HEPES or a sulfate ion in hSC44.20.N32F^L^. Irrespective of the ligand bound to the paratope, the unliganded structures overlay on each other with RMSDs <1.0 Å on Cα carbons of the heavy-chain variable region (Fig. S22*A*). In addition, the apo forms of the antibodies overlay with 3-pTza and 3-pHis bound structures, with minimal deviations of their Cα carbons (Fig. S22*B*). As sulfate and phosphate ions share a tetrahedral geometry, their bonding interactions mimic that of the phosphoryl group from 3-pHis and 3-pTza peptides. Sulfate and phosphate ions make hydrogen bond interactions through the oxygen atoms with the GXGX residues (positions 96-99) in the CDR-H3 loop and Y96^L^ or Y50^H^ (Fig. S22*C,D*). The GXGX motif is also involved in forming hydrogen bonds with HEPES and citrate ions (Fig. 16E,F). Interestingly, the hSC44.20 GXGX motif in CDR-H3 is found to interact with E1^H^ from the N-terminus of the adjacent Fab heavy chain during crystal packing (Fig. S23*A*). E1^H^ also interacts with the GXGX motif in the CDR-H3 loop (Fig. S23*B*). Consequently, the CDR-H1 from the adjacent molecule comes into close contact with the CDR-L1 loop and is displaced from its usual position compared to other structures of hSC44 variants (Fig. S22*A*).

### 3-pHis Specific Antibodies Can Detect Biologically Relevant pHis modifications

To evaluate the sensitivity and utility of the high-affinity, engineered 3-pHis specific antibodies for recognition of biologically relevant pHis proteins, we immunoblotted SDS PAGE-gel separated whole cell lysates from HEK293T cells with hSC44.20 or hSC44.20.N32F^L^ IgGs. These immunoblots were compared with the banding pattern obtained with rSC44. All three antibodies generated similar banding patterns, indicating the presence of 3-pHis signals. This is further confirmed by loss of acid- and thermo-labile pHis signals upon incubation of lysates with acetic acid (pH = 5.5) at 95 °C prior to SDS-PAGE separation (Fig. 5*A*). As rSC44 and its humanized derivatives prefer a GpHAG motif, they predominantly recognize the 3-pHis forms of ACLY (120 kDa) and the α-subunit of SCS (30 kDa) proteins, as observed in the immunoblots with weaker protein signals in other molecular weight ranges (Fig. 5*A*). We then compared the performance of the hSC44.20.N32F^L^ variant in immunofluorescence (IF) staining experiments by co-staining with either rSC44 or rSC1 (1-pTza/pHis specific) antibodies (Fig. 5*B*). hSC44.20.N32F^L^ and rSC44 antibodies concomitantly recognized 3-pHis marks in the cytoplasm of SK-N-BE2 human neuroblastoma cells; as a control, acid-treatment of the slides with 90 °C PBS at pH 4.0 depleted the signals, confirming the specificity of these antibodies for 3-pHis. Notably, we observed 3-pHis localization mainly in the cytoplasm as well as the nucleus, consistent with our previous observations (14). Furthermore, when we co-stained with hSC44.20.N32F^L^ and rSC1 in IF experiments (Fig. 5*B*), hSC44.20.N32F^L^ successfully recognized 3-pHis residues in cells in the cytoplasm, whereas 1-pHis localization was observed primarily in the nucleus. Notably, these co-staining experiments take advantage of the Fc regions from different species (rabbit; rSC1 and human; hSC44.20.N32F^L^) allowing us to probe for 1-pHis and 3-pHis residues in cells simultaneously.

**Fig. 5.**
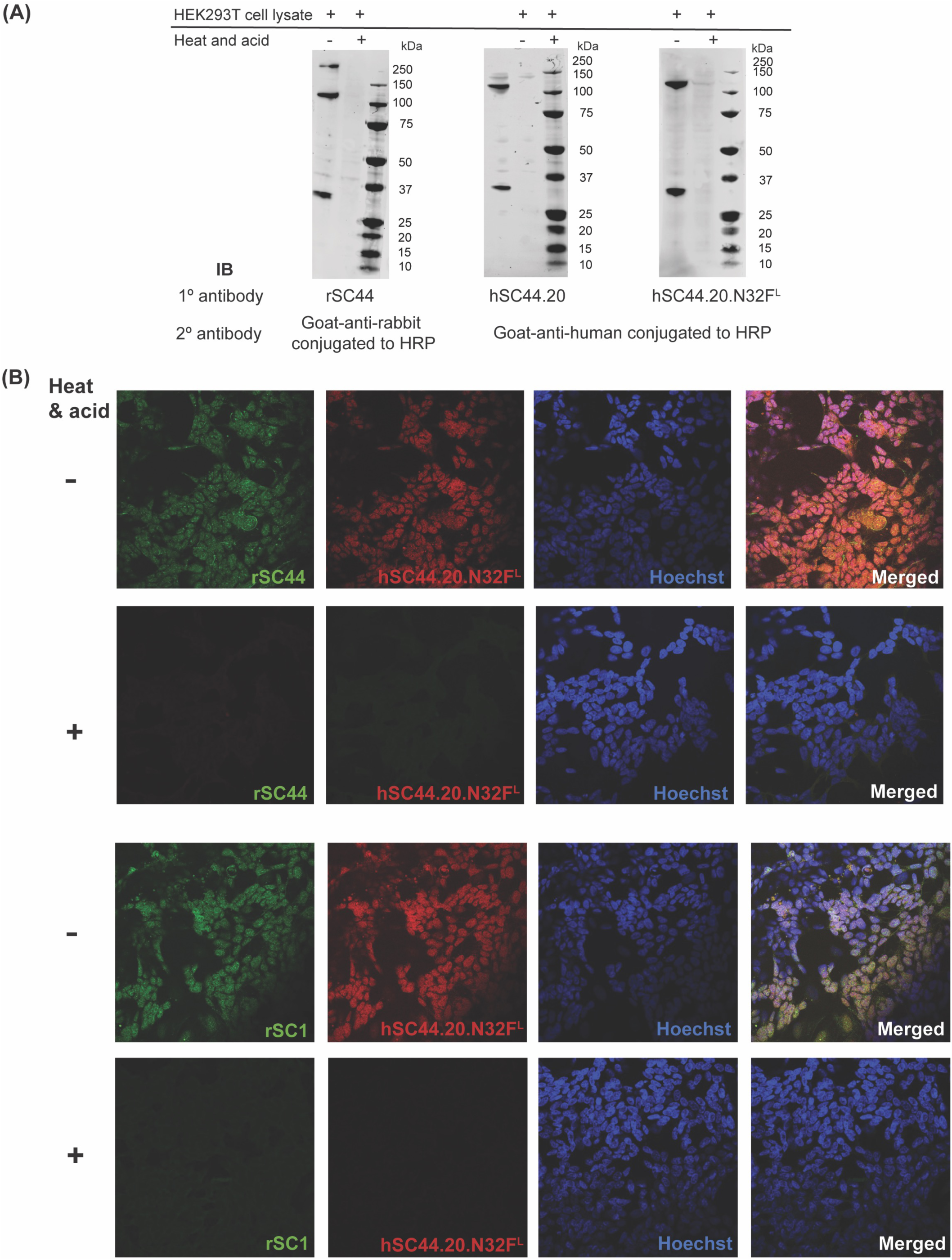
Engineered hSC44 antibodies in typical molecular biology experiments. *(A)* Immunoblot of SDS PAGE-gel separated HEK293T cell lysates probed with rSC44, hSC44.20 and hSC44.20.N32F^L^ IgGs. HRP-conjugated secondary antibodies against rabbit and human IgG were used for detection. Heat and acid treated cell lysates were used as a negative control for pHis signals. *(B)* Immunofluorescence staining of SK-N-BE human neuroblastoma cells probed with rSC44, rSC1 and hSC44.20.N32F^L^ IgGs. Alexa fluor conjugated goat anti-rabbit or goat anti-human antibody was used for immunofluorescence staining. Nuclei were stained with Hoechst and acquired images were merged. Heat and acid treated cells represent negative controls for pHis signals.

## Discussion

The study of pHis in mammalian cells has been historically difficult due to a paucity of tools capable of sensitive detection and quantitative analysis of pHis proteins. The development and use of rabbit antibodies capable of selectively binding 1- and 3-pHis mimetics has enabled new approaches towards understanding the role of pHis in mammalian cell biology (23). Using advanced antibody engineering methods, we were able to solve many of the challenges affecting the development of pHis-specific antibodies from immunized animals. The acid labile and unstable nature of the phosphoramidate bond means that pHis modifications on peptides and proteins are short-lived in animals, and therefore not amenable for use as antigens for the generation of pHis specific antibodies by animal immunization. In these studies, we used prior sequence and structural information of anti-3-pTza rabbit monoclonal antibodies isolated via rabbit immunizations with a 3-pTza peptide to create a humanized version (hSC44) of rSC44 that was amenable to additional engineering. We then constructed phage displayed antibody libraries to engineer hSC44 for improved pHis recognition. Phage display was essential to our strategy, as it allowed us to: (i) control buffer formulations in a pH range that is favorable for pHis stability, (ii) establish a reliable method of 1- and 3-pHis peptide synthesis and isolation, and (iii) implement a carefully controlled selection procedure allowing us to isolate pHis-specific amino acid substitutions in the hSC44 scaffold using a tailored negative and positive selection scheme. Together, this approach coupled with rational engineering of the light chain of the selected variants generated humanized antibodies with favorable developability properties and unparalleled affinity and specificity for 3-pHis. More remarkably, none of the engineered antibodies bound to other phosphorylated amino acids under the conditions used, confirming the 3-pHis specificity of the engineered antibodies. Given the difficulty in generating other phospho-specific antibodies, our work represents a testament to the power of phage display to select antibodies with atomic-level precision.

To complement this approach, we sought to further understand the sets of amino acid substitutions in the hSC44 CDRs that governed high affinity and selectivity for 3-pHis. To do this, we undertook an extensive crystallization campaign and now report for the first time several structures of antibodies complexed with pHis peptides. To aid our structural studies, we used novel technologies that favored antibody crystallization by promoting interactions at interfaces away from the paratope (32–34). These substitutions are located in the elbow region of the heavy chain as well as the constant region of the kappa light chain and enabled us to obtain crystals that lead to 11 structures consisting of hSC44, hSC44.20 and hSC44.20.N32F^L^ in the apo form as well as in complex with 3-pTza and 3-pHis peptides. This large dataset revealed the molecular determinants governing high affinity interactions with 3-pTza/3-pHis peptides in hSC44 variants. Additionally, QM/MM analysis offers a molecular-level insight into the effect of the mutation (N32F^L^), which leads to a newly established pi-pi interaction with pHis, resulting in improved binding affinity to pHis by hSC44.20.N32F^L^ in comparison to hSC44.20.

The high affinity of isolated antibodies for 3-pHis was likely crucial for the successful crystallization of antibodies in complex with a 3-pHis phosphorylated peptide, possibly by preventing the spontaneous and rapid hydrolysis of the pHis moiety (24). The comparison between 3-pTza and 3-pHis peptide-bound hSC44 variants demonstrated that CDRs of these variants interact with the modifications in a similar way. However, the large affinity difference between the 3-pTza versus 3-pHis bound antibodies (>1000 fold) could not be explained by the crystal structures. Hence, we resorted to QM/MM analysis of the structures which revealed that the imidazole ring makes the phosphoryl moiety less electronegative compared to the triazolyl ring by drawing electrons from the phosphoryl moiety, thereby decreasing the number of interactions with the paratope. Another positive feature of hSC44 is its absolute isomer specificity, as we did not observe any interaction with 1-pTza or 1-pHis peptides using BLI. This property was further confirmed during crystallization, as the native peptides were chemically phosphorylated using a strategy that results in a mixture of unphosphorylated, 1-pHis, 3-pHis and 1,3-di-pHis peptides (see Materials and Methods for a detailed description). We subjected this mixture to co-crystallization with hSC44 variants without peptide purification, and in every case, we obtained crystals of antibodies in complex with 3-pHis peptides. We failed to obtain a purely apo structure of any of the hSC44 variants, as the positive electrostatic potential of its CDRs facilitated interactions with small electronegative anions from either the purification buffers, crystallization buffers, or neighboring Fabs. However, this is likely an artifact of crystallization and the high concentration of anions, as we did not observe binding to other negatively charged molecules (i.e., ssDNA, dsDNA) in our specificity ELISA (Fig. S8) nor to other phosphoryl-based PTMs such as pSer, pThr, pTyr by ELISA (Fig S5*D*) or the 1-pHis isoform (Tables 1-2), as measured by BLI. Overall, our structural analysis of these Fabs reveals that humanization and engineering did not alter the hallmark properties of the original rabbit rSC44-8 antibody. These antibodies recognize the phosphoryl moiety through a GXGX motif (‘nest’ motif) that was retained during engineering. They displayed properties of isomer specificity and structures also reveal that they retained the property of sequence dependence towards GpHAGA motif in the epitope. However, individual changes at the single residue level in hSC44.20.N32F^L^ contribute to its higher affinity binding to 3-pHis.

There are several important advances made by the work presented here. First, we developed a workflow that allows the use of unstable pHis for *in vitro* engineering of anti-3-pHis antibodies. This *in vitro* selection protocol can be expanded to other non-canonical and unstable PTMs. There are 9 amino acids with modifiable side chains and six, like pHis, are understudied due to a lack of reagents for their analysis (23). We anticipate that the methods developed in this study can be adapted for isolating antibodies recognizing additional non-canonical PTMs. Second, the high affinity 3-pHis specific antibodies described here can be used to probe the pHis phosphoproteome in greater depth using immunoblotting, immunofluorescence staining, and phosphoproteomics. To complement these tools, we have generated an unprecedented amount of structural data on pHis recognition. These data, and particularly the N32F substitution, can be exploited to construct next-generation phage-displayed libraries of pHis specific Fabs and to rationally engineer sequence-specific pHis antibodies. In conclusion, our work demonstrates how *in vitro* evolution of PTM-specific antibodies can be used to generate a novel set of tools for investigating the biology of pHis.

## Materials and Methods

### IgG/Fab Protein Expression and Purification

IgG/Fab protein was expressed as previously described (47) with some modifications. Briefly, plasmids designed to express the heavy and light chains were co-transfected into Expi293 cells (ThermoFisher Scientific; A14527) using the FectoPRO^®^ transfection reagent kit (Polyplus) according to manufacturer’s instructions. After 5 days of expression, cell culture media was harvested and applied to CaptoL affinity column (Cytiva; 17547806). IgG/Fab protein was eluted using IgG elution buffer (Pierce™; 21009) and neutralized with 1 M Tris-HCl (pH = 8.0). Fractions containing IgG/Fab protein were combined, concentrated, and buffer exchanged into 1x HBS (50mM HEPES (pH 8.5), 150 mM NaCl) using Amicon Ultra-15 centrifugal units (50 kDa/10 kDa respectively, Millipore Sigma; UFC9010). IgG/Fab protein was characterized for purity by SDS-PAGE with concentrations being determined by spectrophotometry (280 nm).

For crystallization, Fabs were expressed as described, but were affinity purified from the cell culture media with CaptureSelect CH1-XL resin (Thermo-Fisher) followed by a wash with PBS and elution with 50 mM acetate, pH = 4.0, and neutralization with 1 M Tris-HCl (pH = 9.0).

### Size Exclusion Chromatography

IgG/Fab protein was purified by size exclusion chromatography (SEC) using an ÄKTA system (GE Healthcare) equipped with a Hi-Load 16/60 Superdex 75 gel filtration column (GE Healthcare Life Sciences). SEC columns were pre-equilibrated and run with 1x HBS with a flow rate of 1 mL/min. Fractions containing IgG/Fab protein were combined and concentrated using Amicon Ultra-15 centrifugal units (50 kDa/10 kDa, respectively, Millipore Sigma; UFC9010). IgG/Fab protein was characterized for purity by SDS-PAGE with concentrations being determined using spectrophotometry (280 nm). Fabs for crystallization were purified by SEC with running buffer consisting of 20 mM Tris and 150 mM NaCl.

### Peptide Synthesis

Phosphohistidine mimetic (1/3-pTza) peptides (Biotin-AminocaproicAcid-A.G.A.G.1-pTza.A.G.A.G; 1-pTza peptide and Biotin-AminocaproicAcid-A.G.A.G.3-pTza.A.G.A.G; 3-pTza peptide) and histidine containing peptide (Biotin-AminocaproicAcid-A.G.A.G.H.A.G.A.G; His-peptide) were synthesized as previously described (23).

Peptides ACLYana-3-pTza and ACLYana used in the crystallization experiments were synthesized by the Salk Peptide Synthesis Core. Both peptides have the same sequence Ala-Gly-Ala-Gly-X-Ala-Gly-Ala-Gly except for one residue. X is 3-pTza (3-phosphotriazolyl alanine) in ACLYana-3-pTza peptide and histidine in ACLYana peptide. These peptides have an acetyl group at the N-terminus and an amide at the C-terminus. They were synthesized using standard Fmoc chemistry on a Gyros Protein Technologies Tribute peptide synthesizer with UV monitoring. After deprotection, peptides were analyzed by mass spectrometry and analytical HPLC by the Proteomics Core at the Salk Research Institute. The peptides were purified by HPLC to 99% purity by 21^st^ Century Biochemicals, Inc. The Fmoc-pTza molecules were synthesized at 10 g scale for incorporation into the peptides.

### Preparation of rProtein A-IgG Matrix

A 100 µL aliquot of rProtein A Sepharose® beads (Cytiva; 17-1279-01) was washed with 1 mL dH_2_O three times and equilibrated in 1 mL 1x HBS. 100 µg rSC1 (1-pTza/pHis) IgG (24) was added to one-half of equilibrated rProtein A beads and 100 µg of rSC44 (3-pTza/pHis) (24) IgG to the other half of equilibrated rProtein A beads, to make two separate resins. IgG binding to rProtein A beads was carried out at 4 °C for 1 h with end-over-end rotation. rProtein A-IgG resin mixture was then spun down at 1000 rpm for 1 min, supernatant was removed, and the beads were washed with 1 mL 1x HBS to remove unbound IgG. This was repeated three times. IgGs were then crosslinked to rProtein A resin by adding BS^3^ (Thermo Scientific™; 21580; resuspended in 20 mM HEPES, pH = 8.5) to a final concentration of 1 mM. The IgG-rProtein A-BS^3^ mixture was then incubated at room temperature for 1 h with end-over-end rotation. The reaction was quenched by adding an equal volume 1M Tris-HCl (pH 8.5) and incubated at room temperature for 10 mins with end-over-end rotation. The resin was spun down at 1000 rpm for 1 min, supernatant was removed, and the resin was washed with 1 mL 1x HBS to remove unbound BS^3^. This was repeated 3 times and resin was stored in 300 µL 1x HBS with 0.05% NaN_3_ at 4°C.

### Phosphoramidate Reaction and Phosphohistidine Peptide Production

The synthesis of potassium phosphoramidate and the phosphorylation of His side-chains using potassium phosphoramidate have been described previously (48). Briefly, 10 mg of potassium phosphoramidate was resuspended in 1 mL of 1x HBS (10 µg/µL). 100 µg of His-peptide (biotin-aminocaproic-acid.A.G.A.G.H.A.G.A.G) resuspended in 1x HBS was mixed with 1 mg phosphoramidate (100 µL, 1:10 ratio). Reaction was incubated overnight at room temperature for 16 h and subsequently quenched with an equal volume of 1 M Tris-HCl (pH 8.5).

### Phosphohistidine Peptide Immunoprecipitation

Preparation of rProtein A-IgG resins. 1-pHis and 3-pHis resins were washed with 1 mL 1x HBS three times to remove NaN_3_. Half of the His-peptide:potassium phosphoramidate reaction was added to anti-1-pHis mAb (SC1-1) resin and the other half to anti-3-pHis (SC44-8) mAb resin and incubated at room temperature for 1 h with end-over-end rotation. Resins were washed thrice with 500 µL 1x HBS by centrifugation at 1000 RPM. Bound pHis peptides were eluted by adding 500 µL 2 M Imidazole (re-suspended in 1x HBS, pH 8.5) to resin and incubating at room temperature with end-over-end rotation for 10 min. This was repeated once more for a total elution volume of 1 mL. Elution volume can change depending on desired use or concentration of peptide. Typical yields using this protocol were ∼500 ng of phosphopeptide and can be scaled up as needed. Peptide concentrations were estimated using a quantitative colorimetric peptide assay kit (Pierce™; 23275) according to manufacturer’s instructions. We also developed a quantitative ELISA assay using 3-pTza peptides and r/hSC44 to generate standard curves for the estimation of pHis peptide concentrations.

Regeneration of rProtein A-IgG resins. Used resins were washed with 1 mL dH_2_O three times. To strip any 1/3-pHis peptide still bound to the resin, 1 mL 50 mM sodium acetate, pH = 4.0 was added and incubated at room temperature for 10 min with end-over-end rotation. This was repeated two times. Resins were then washed with 1 mL 1x HBS three times and resuspended in 300 µL 1x HBS with 0.05% NaN_3_ and stored at 4 °C.

### ELISA

A Nunc® MAXISORP™ 384-well ELISA plate (Millipore Sigma; P6366) was coated with 25 µL/well of 2 µg/µL streptavidin (New England Biolabs; N7021S) diluted in 1x HBS and incubated overnight at 4 °C. For experiments involving phage-ELISAs, anti-FLAG M2 mouse derived IgG (MilliporeSigma; F3165) was diluted 1/5000 in 1x HBS and 25 µL/well was dispensed to appropriate wells and incubated overnight at 4°C. The following day, plates were washed four times with 1x HBS-T (1x HBS, 0.05% Tween 20) with 90 µL/well. Plates were then blocked with 75 µL/well 1x HBS, 0.5% BSA for 1 h at room temperature with shaking, followed by plate washing. Biotinylated peptides were added to the plate at various concentrations at 25 µL/well, incubated at room temperature for 30 mins with shaking, followed by plate washing. IgG was added at various concentrations at 25 µL/well and incubated at room temperature for 1 h with shaking, followed by plate washing. For the detection of rabbit IgG (rSC44), HRP conjugated goat anti-rabbit IgG (Thermo Scientific™/Invitrogen; A16110) diluted 1/5000 in 1x HBS-T, 0.05% BSA was added at 25 µL/well and incubated at room temperature for 30 mins with shaking, followed by plate washing. For humanized IgGs (hSC44 variants), HRP-conjugated goat anti-Human Kappa (SouthernBiotech; 2060-05) was used with identical conditions. For Fab-phage ELISA (hSC44 variants), HRP conjugated mouse anti-M13 (Sino Biological; 11973-MM05T-H) was used at a 1:12000 dilution. Binding was detected using 25 µL/well 3,3′,5,5′-tetramethylbenzidine (TMB) (Thermo Scientific™; 34021) chromogenic substrate and reactions were quenched using 25 µL/well 1 M phosphoric acid. Plate absorbance values (450nm) were read using a PowerWave XS microplate reader (BioTek).

For quantitative ELISA to estimate 3-pHis peptide concentration, a 2-fold serial dilution (12 dilutions total) of 1/3-pTza peptides starting at ∼3.5 mM (∼4 mg/mL) was added to streptavidin coated plates. A 10-fold serial dilution (6 dilutions total) of purified biotinylated 3-pHis peptide was also added to separate wells on streptavidin coated plates. Wells were probed with 100 nM rSC44 and ELISA was carried out as previously described. Standard curves corresponding to linear detection range of 3-pTza peptides were generated using Prism 6 (GraphPad Prism) and concentrations of 3-pHis with absorbance values falling within the linear range were solved. Concentrations of 3-pHis peptides were extrapolated back to undiluted sample and averaged. Three technical replicates were conducted for each measurement.

For EC_50_ measurements using hSC44 variant IgGs, 100 nM peptide was added to each streptavidin coated well. A 10-fold serial dilution (7 dilutions total) of IgG starting at 1 µM was added to the plate. Streptavidin and 1x HBS were used as negative controls and rSC44 was used as a positive control. ELISA was carried out as previously described and EC_50_ curves were calculated (Graphpad Prism). Experiments were performed with 4 technical replicates and 3 experimental replicates.

For phage ELISA using hSC44 variant Fab displayed on phage after selections, 1 µM peptide was added to each streptavidin coated wells. While streptavidin and 1x HBS were used as negative controls and rSC44 was used as a positive control. hSC44 phage pools were first normalized based on FLAG expression using anti-FLAG M2 ELISA. Briefly, a 2-fold serial dilution (7 dilutions total) of overnight phage pools was added to the wells of a plate coated with anti-FLAG M2 antibody (Sigma). hSC44 variant concentrations were then normalized based on FLAG display and used to probe peptide containing plates. Experiments were performed with 4 technical replicates and 3 experimental replicates.

For cross-reactive ELISAs, 25 µL/well of select macromolecules were added to wells at final concentrations of cardiolipin (50 µg/mL; C0563; Sigma), keyhole limpet hemocyanin (KLH; 5 µg/mL, H8283; Sigma), lipopolysaccharide (LPS; 10 µg/mL; tlrl-eblps; InvivoGen), single-stranded DNA (ssDNA; 1 µg/mL; D8899; Sigma), double-stranded DNA (dsDNA; 1 µg/mL; D4522; Sigma), insulin (1 µg/mL; 91077C; Millipore Sigma), BSA (1% (w/v)), streptavidin (10 µg/mL) and goat anti-human Fc (1 µg/mL; 109-005-098; Jackson). IgGs being tested for cross-reactivity were added at 100 nM in a volume of 25 µL/well. ELISAs were then carried out as previously described.

### Construction of Phage-Displayed Libraries

Phage-displayed library construction methods fusing Fab to phage coat protein III (pIII) of M13 bacteriophage have been previously described (49). Here we use some minor modifications for obtaining hSC44 phage-displayed libraries. Briefly, residues that were oriented towards, proximal to (within 10 Å) or directly contacting the 3-pTza peptide ligand in a crystal structure (PDB ID: 6X1V) were selected for mutagenesis for libraries hSC44 1-5 (Fig S2-3). Key framework residues required for CDR orientation were selected for mutagenesis for library hSC44 6. Appropriately designed “stop template” versions of each proposed library were synthesized by introducing stop codons into regions intended to be mutagenized. Mutagenic oligonucleotides targeting these positions while also repairing stop codons were synthesized (Integrated DNA Technologies) and used to construct libraries (Table S1). Mutagenic oligonucleotides were synthesized to adjust the nucleotide ratio of diversified positions to 70% of the wild-type nucleotide and 10% of each of the other nucleotides retaining the wild-type amino acid roughly 50% of the time (42). The libraries diversities ranged from 10^9^-10^10^ unique variants.

### Affinity Selections

Libraries hSC44 1-6 were pooled and cycled through five rounds of affinity selections on 96-well Maxisorp Immunoplates (Nunc™ Thermo Scientific™; 12-565-135) as previously described (25), with some modifications. Selections were carried out as outlined in Fig 2*A*. 3-pHis peptides used in selections were generated fresh each day (see Phosphoramidate Reaction and Phosphohistidine Peptide Production section of Materials and Methods). Plates were coated with 100 µL/well of reagent indicated in Table S2 overnight at 4 °C and blocked the following day with 1x HBS containing 0.5% BSA. Phage pools were cycled through negative and positive selections by incubation at room temperature for 1 hour with shaking for each condition. Washing steps after positive selection used 1x HBS-T, 0.05% Tween-20 with 2-5 min incubations with shaking at each washing step. Phage were eluted from positive selection using 100 µL/well 100 mM HCl with incubation for 10 min with shaking at room temperature, eluted phage pools were neutralized with 1 M Tris-HCl (pH 11) and used to infect OmniMax™ (Invitrogen™; C854003) *Escherichia coli (E. coli)* cells at OD = 0.6-0.8 for overnight amplification of phage pools.

### Biolayer Interferometry

Binding kinetics and estimation of affinity (K_D_) measurements were carried out using an Octet HTX instrument (FortèBio) as previously described (50) with some minor modifications. Biotinylated 1/3-pTza/pHis peptides and non-phosphorylated His-peptides were captured on streptavidin biosensors at 20 nM. Assay buffer used was 1x HBS-T, 1% BSA. IgGs were used at various concentrations (20-200 nM).

### Data Analysis

ELISA data were analyzed using Prism 6 software (GraphPad Software Inc). Structural alignments were processed and visualized in PyMol (v.2.5.2). Library design and DNA sequencing was analyzed using Geneious v.10.0.09 (https://www.geneious.com). Biolayer interferometry data were analyzed using Octet Systems Software v.9.0 (FortèBio).

### Crystallization and Structure Solution

ACLYana-pHis peptides were prepared by incubating 50 mM of unphosphorylated ACLYana peptide with 250 mM of phosphoramidate in HBS for 12 h at room temperature. The resulting reaction has mixed species of unphosphorylated peptide, phosphorylated peptide with 1-pHis modification and 3-pHis modification. The whole reaction was used for co-crystallization. Fab versions of hSC44, hSC44.20 and hSC44.20.N32F^L^ antibodies described in Table S4-5 were pre-incubated with either ACLYana-3-pTza or ACLYana-pHis peptides at a molar ratio of 1:5 (Fab:peptide). hSC44 Fab variants (hSC44, hSC44.20 and hSC44.20.N32F^L^) complexed with ACLYana-3-pTza peptides as well as unliganded hSC44 proteins were screened for crystallization using our Rigaku CrystalMation system at Scripps Research, with JCSG Screens 1-4 at 4 °C and 20 °C. The crystallization conditions used are reported in Table S5. Data were collected at either APS beamline 23-ID-D, SSRL beamline 12-2 or ALS beamline 5.0.3 at 100 K. The diffraction images were indexed, integrated, and scaled using either HKL-2000 (51) or XDS (52). The rSC44 Fab (PDB ID: 6X1V) (24) was used as search model for molecular replacement with PHASER-MR (53) to obtain initial phase information. Refinement was carried out by alternating rounds of Phenix.refine (54) and model building software Coot (55). The final models were validated using MolProbity (56) and figures were prepared using PyMOL (Schrodinger LLC) and UCSF Chimera (57). Buried molecular surfaces were analyzed with the MS program (58) using a 1.7 Å probe radius. The pi-pi interaction distances and angles were measured using either BINANA 2.0 or PLIP (59). The What If server was used to measure torsion angles (60). The x-ray data collection, refinement statistics and the model validation values are reported in Table S4.

### Quantum Mechanics/Molecular Mechanics (QM/MM)

The experimentally resolved structures (hSC44.20:ACLYana-3-pTza, hSC44.20:ACLYana-3-pHis and hSC44.20.N32F^L^:ACLYana-3-pHis) underwent QM/MM minimizations to achieve geometry-optimized structures of the active site at the ab initio DFT level, allowing assessment of the quantitative response of the ligand-bound structures to mutations. The MM region was modelled using the OPLS force field [3], while the active site was treated with the LACVP* basis sets, combining the successful 6-31G basis set with the LANL2DZ effective core basis set (61, 62) as implemented in the Schrodinger Qsite tool (63–65). QM/MM calculations were performed in a non-periodic manner, with atoms more than 30 Å from the ligand (antibody: pHis/pTza) excluded, and those between 20 and 30 Å fixed. Population and Orbital analyses for the QM region were conducted using the Jaguar tool (66) as implemented in Schrodinger 2023-2. Additionally, QM analysis was also performed on the antibodies (pHis and pTza) using the Jaguar tool (66), as implemented in Schrodinger-2023. The peptides are locally aligned to analyze the shift in the antibody near the Fab region caused by the N32F mutations using UCSF Chimera (57). For graphics preparation and visualization, we used UCSF Chimera and Maestro-2023 from the Schrodinger suite tool.

### Molecular Dynamics and Binding Affinity (FEP) Simulation Details

For molecular dynamics-based contact analysis and FEP simulations, the starting model was the experimentally resolved structures (hSC44.20:ACLYana-3-pTza, hSC44.20:ACLYana-3-pHis and hSC44.20.N32F^L^:ACLYana-3-pHis).

### Contact Analysis

MD-based contact analysis was performed on three experimentally resolved structures (hSC44.20:ACLYana-3-pTza, hSC44.20:ACLYana-3-pHis and hSC44.20.N32F^L^:ACLYana-3-pHis) using the Maestro-Desmond Interoperability Tools (67) in Schrodinger. Each system was solvated with ∼40,000 TIP3P water molecules, depending on the antibody and mutation. Water molecules directly involved in metal coordination were retained from the original starting structure. All three systems were prepared using the Protein Preparation Wizard in Schrodinger. Minimization was conducted using a steepest descent algorithm for 100,000 steps, followed by a 20 ns equilibration with restrained heavy atoms (backbone heavy atoms of the protein and nucleic acids with an isotropic force of 1000 kJ mol⁻¹nm⁻¹) under constant number, pressure, and temperature (NPT) and constant number, volume, and temperature (NVT) conditions up to 1 ns at 300 K using the standard MD procedure in the Desmond tool of Schrodinger 2023.

### Criteria of Contact Analysis

The contacts presented here are chosen based on these criteria.

### H-bonds

The current geometric criteria for protein-ligand H-bond are: (1) distance of 2.5 Å between the donor and acceptor atoms. (2) (D—H···A); A donor angle of 120°-180° between the donor-hydrogen-acceptor atoms (D—H···A). (3) An acceptor angle of 120°-180° between the hydrogen-acceptor-bonded atom atoms (H···A—X).

### Water Bridge

Hydrogen-bonded protein-ligand interactions mediated by a water molecule. The current geometric criteria for a protein-water or water-ligand H-bond are: (1) A distance of 2.8 Å between the donor and acceptor atoms (D—H···A); a donor angle of 120°-180° between the donor-hydrogen-acceptor atoms (D—H···A); (2) An acceptor angle of 120°-180° between the hydrogen-acceptor-bonded atom atoms (H···A—X).

### Hydrophobic Contacts

The current geometric criteria for hydrophobic interactions are as follows: (1) π-Cation — Aromatic and charged groups within 4.5 Å. (2) π-π — Two aromatic groups stacked face-to-face or face-to-edge within 3.5-5 Å.

### Binding Affinity (FEP) Simulations

The FEP+ program (68, 69) from the Schrödinger suite 2023-2 was used to determine the relative binding free energies of pHis between the wild-type and mutant N32F, evaluating the effect of the N32F mutation on pHis binding. The wild-type and mutant complexes were prepared using the Protein Preparation Wizard. For each wild-type complex and its corresponding mutant, the FEP Protein Mutation for Ligand Selectivity GUI was used to build a perturbation map. The OPLS4 force field modelled proteins and ligands (70) and torsion parameters for all ligand fragments were checked using a Force Field Builder. A 10 Å cubic box filled with approximately ∼ 40,000 TIP3P water molecules (71) was used for the complex and solvent perturbation leg (wild-type and mutant systems). The number of alchemical λ windows was set to 12 by default to connect the wild-type and mutant states. For each λ window, production MD was run for 20 ns in the NPT ensemble. The Bennett acceptance ratio method (72) (BAR) was used to calculate the free energy differences. Mutational free energy calculations were performed in triplicate, initializing the MD with different random seeds. The reported relative binding free energies were calculated as the average of the three ΔΔG values from independent simulations.

### Immunoblotting

The protocol used here was described previously (73). To maintain pHis signals, cell lysates were processed at 4 °C and alkaline buffer conditions were used throughout the experiment. HEK293T cells were grown in DMEM medium in a 10-cm dish for 3 days before processing. Cells were washed twice with ice-cold 1x PBS buffer, detached from the dish surface using a cell scraper and lysed with 400 µL of lysis buffer (25 mM Tris pH 8.5, 140 mM NaCl, 0.1 % Tween-20, 1 mM PMSF, 1x protease inhibitor tablet, and 1x PhosStop tablet. Cells were lysed by sonicating for 1 min using a 10 sec on and 10 sec off cycle at 40% amplitude. Lysates were clarified by centrifugation at 12,000 rpm at 4 °C and 30 µg of supernatant was loaded onto an alkaline 12 % SDS-PAGE gel followed by alkaline immunoblotting as previously described (Methods Kalagiri Ref). Electrophoresis and immunoblotting were carried out 4 °C. Lysates treated with acid (pH 5.5) and heat (95 °C for 15 min) were used as negative controls. Membranes were blocked with blocking buffer (5 mM Tris pH 8.5, 30 mM NaCl, 0.1 % Tween-20, and 0.1 % casein) and then incubated with 0.5 µg/mL of primary antibodies (rSC44, hSC44.20, or hSC44.20.N32F^L^) diluted in blocking buffer for 1 h at room temperature. Membranes were washed three times with wash buffer (25 mM Tris pH 8.5, 140 mM NaCl, 0.1 % Tween-20) with 10 min incubation times. The blot was then incubated for 1 h at room temperature with 0.5 µg/mL of goat anti-human kappa antibody conjugated to HRP (31460, SouthernBiotech) if hSC44.20, or hSC44.20.N32F^L^ were used as primary antibodies and goat anti-rabbit antibody conjugated to HRP (31460, Thermo Fisher) if rSC44 was used as a primary antibody. Membranes were washed with wash buffer three times and developed using luminol reagent which is a substrate for HRP. The resultant chemiluminescence was detected using BioRad ChemiDoc MP Touch Imaging System.

### Immunofluorescence

**SKN-BE**(2) cells were fixed with 4% paraformaldehyde (pH 8.5) for 15 min at room temperature and washed three times with 4 °C PBS (pH 8.5) (control slides and coverslips were washed three times with 90 °C PBS at pH 4.0). Then, the cells were permeabilized with 0.2% Triton X-100 for 15 min at room temperature and washed three times with 4 °C PBS (pH 8.5) (control slides and coverslips were washed with 90 °C PBS at pH 4.0). Cells were blocked in 10% BSA in TBS-T (pH = 8.5) at room temperature for 30 min, before incubating with primary antibody diluted in 1% BSA in TBS-T (pH 8.5) at 4°C overnight. The dilutions were used as follows: 2 µg/mL for rSC44-8 and rSC1-1 and 8 µg/mL for hSC44.20.N32F^L^. The day after, cells were washed 3 times for 5 minutes with PBS at room temperature. Cells were then incubated with secondary antibodies diluted in 1% BSA in TBS-T (pH 8.5) for one hour as follows: 1/200 for secondary goat anti-rabbit IgG antibody Alexa Fluor® 488 (A-11034, Invitrogen, Carlsbad, CA, USA), 1/20 for secondary mouse anti-human Ig light chain K antibody Alexa Fluor® 647 (316514, BioLegend®, San Diego, CA, USA) and mixed with Hoescht 33342 (H3570, Invitrogen, Carlsbad, CA, USA). Finally, the cells were washed 3 times with PBS (pH 8.5) at room temperature and then mounted with Prolong Gold Antifade (P36930, Invitrogen, Carlsbad, CA, USA). Cell imaging was performed with an SP8 confocal microscope (Leica) using a 63×/NA 1.4 Plan Neofluor objective lens. Each channel was imaged sequentially using the multitrack recording module before merging in order to prevent contamination between fluorochromes.

## Acknowledgments

GM/CA@APS has been funded by the National Cancer Institute (ACB-12002) and the National Institute of General Medical Sciences (AGM-12006, P30GM138396). This research used resources of the Advanced Photon Source; a U.S. Department of Energy (DOE) Office of Science User Facility operated for the DOE Office of Science by Argonne National Laboratory under Contract No. DE-AC02-06CH11357. The Eiger 16M detector at GM/CA-XSD was funded by NIH grant S10 OD012289. Use of the Stanford Synchrotron Radiation Lightsource, SLAC National Accelerator Laboratory, is supported by the U.S. Department of Energy, Office of Science, Office of Basic Energy Sciences under Contract No. DE-AC02-76SF00515. The SSRL Structural Molecular Biology Program is supported by the DOE Office of Biological and Environmental Research, and by the National Institutes of Health, National Institute of General Medical Sciences (including P41GM103393). The contents of this publication are solely the responsibility of the authors and do not necessarily represent the official views of NIGMS or NIH. The Berkeley Center for Structural Biology is supported in part by the Howard Hughes Medical Institute. The Advanced Light Source is a Department of Energy Office of Science User Facility under Contract No. DE-AC02-05CH11231. The Pilatus detector on 5.0.1. was funded under NIH grant S10OD021832. The ALS-ENABLE beamlines are supported in part by the National Institutes of Health, National Institute of General Medical Sciences, grant P30 GM124169. R.M.L was supported by NIH R35 GM132090 and U01 AI136680. D.L was supported by the Hearst Foundations grant U01 AI136680. A.B. would like to acknowledge support from the Eric and Wendy Schmidt AI in Science Postdoctoral Fellowship, a program of Schmidt Sciences. The Salk Peptide Synthesis and Proteomics Cores are supported by an NCI-designated Cancer Center Support Grant P30 CA014195. T.H. is a Frank and Else Schilling American Cancer Society Professor and holds the Renato Dulbecco Chair in Cancer Research. T.H. is supported by an NIH National Cancer Institute R35 (5 R35 CA242443) award, R.K. was supported by a Salk Innovation Award and Salk Alumni Fellowship, and M.S. was supported by a Hewitt Foundation Fellowship. The structure work was supported in part by the Skaggs Institute of Chemical Biology (I.A.W).

## Data deposition

The X-ray coordinates and structure factors of the pHis Fab:peptide complexes and pHis Fabs have been deposited in the Protein Data Bank with accession codes-8UJI (hSC44.S1C:AGAG-3pTza-AGAG); 8UIT (hSC44.S1C:AGAG-3pHis-AGAG); 8UIO (hSC44.S1C); 8UIH (hSC44.S1C.20:AGAG-3pTza-AGAG); 8UIG (hSC44.S1C.20:AGAG-3pHis-AGAG); 8UHT (hSC44.S1C.20); 8UHS (hSC44.S1C.20); 8UHP (hSC44.S1C.20.N32F^L^:AGAG-3pTza-AGAG); 8UHN (hSC44.S1C.20.N32F^L^:AGAG-3pHis-AGAG); 8UHJ (hSC44.S1C.20.N32F^L^); and 8UHH (hSC44.S1C.20.N32F^L^).

