## Supplementary material for "Using phage display for rational engineering of a higher affinity humanized 3’phosphohistidine-specific antibody"

Fig. S1.

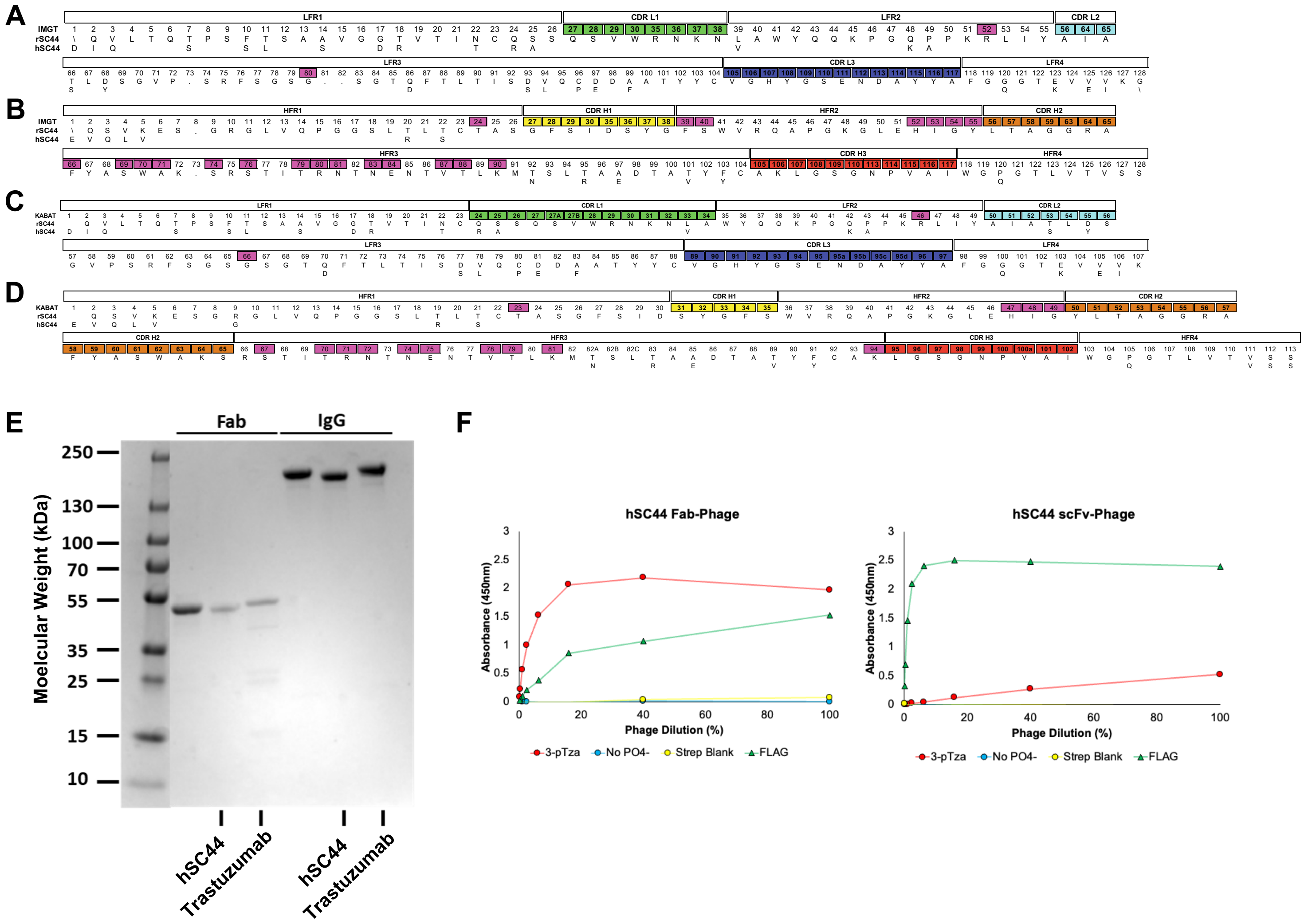
A
B
C
D
E
Fab
IgG
Moelcular Weight (kDa)
250
130
100
70
55
35
25
15
10
hSC44
Trastuzumab
hSC44
Trastuzumab
F
hSC44 Fab-Phage
hSC44 scFv-Phage
Absorbance (450nm)
Phage Dilution (%)
3-pTza
No PO4-
Strep Blank
FLAG

**Fig. S2.**

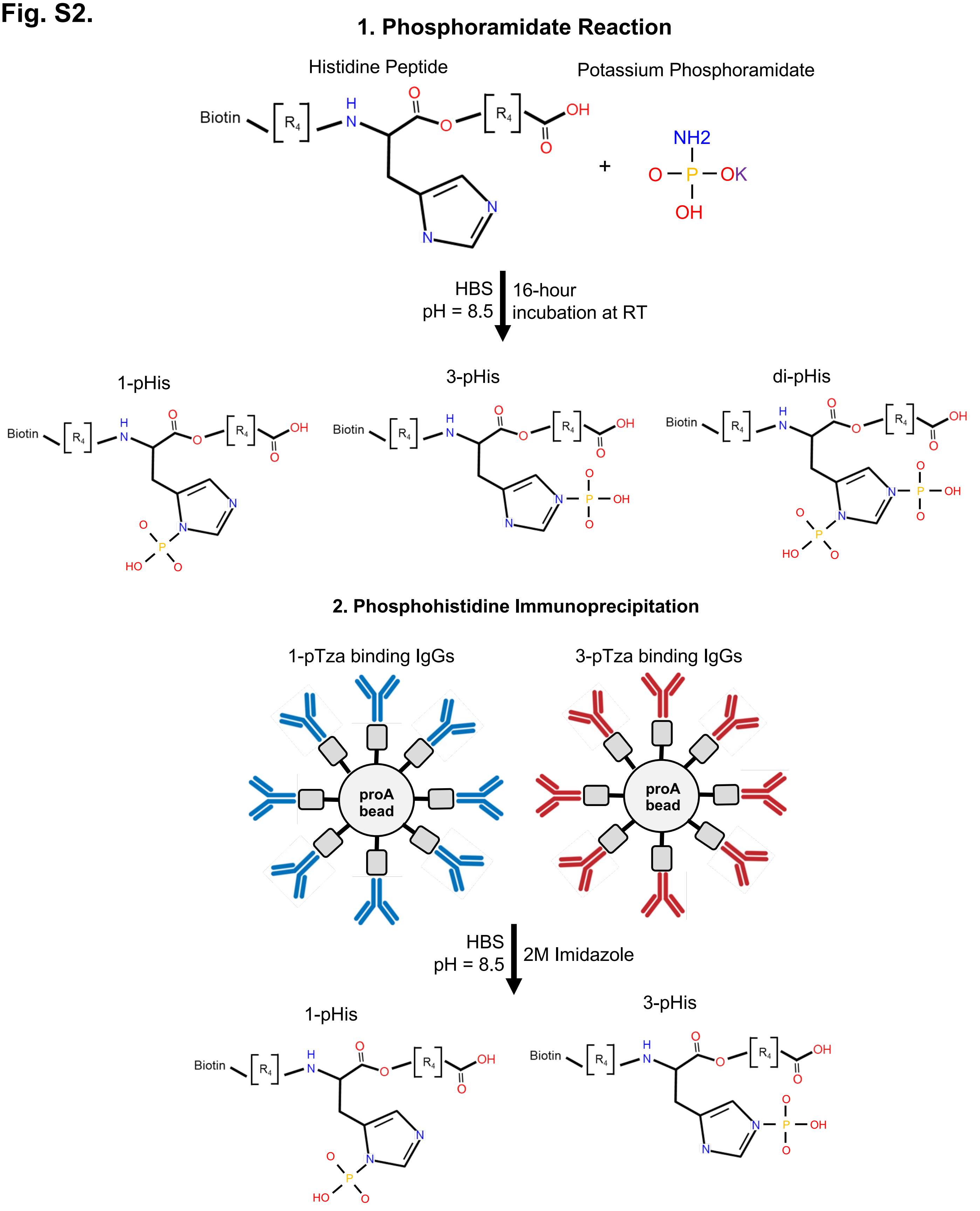

1. Phosphoramidate Reaction
Histidine Peptide
Potassium Phosphoramidate
Biotin
R4
H
N
O
OH
+
NH2
P
OK
HBS
pH = 8.5
16-hour
incubation at RT
1-pHis
3-pHis
di-pHis
HO
2. Phosphohistidine Immunoprecipitation
1-pTza binding IgGs
3-pTza binding IgGs
proA
bead
HBS
pH = 8.5
2M Imidazole
1-pHis
3-pHis

**Fig. S3.**

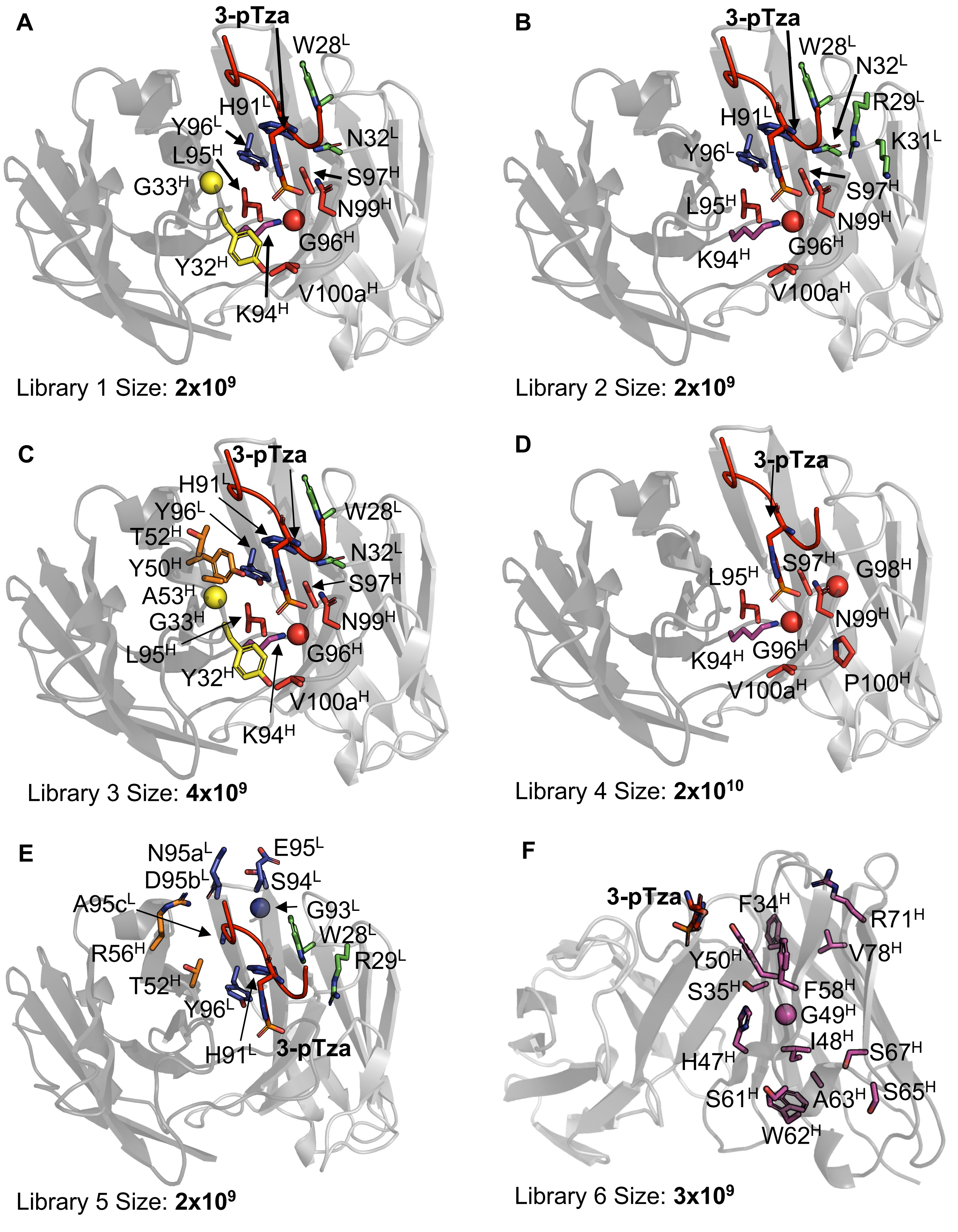
A
3-pTza
W28$^{L}$
H91$^{L}$
Y96$^{L}$
N32$^{L}$
L95$^{H}$
S97$^{H}$
G33$^{H}$
N99$^{H}$
G96$^{H}$
Y32$^{H}$
V100a$^{H}$
K94$^{H}$
Library 1 Size: **2x10$^{9}$**
B
3-pTza
W28$^{L}$
N32$^{L}$
R29$^{L}$
H91$^{L}$
K31$^{L}$
Y96$^{L}$
S97$^{H}$
L95$^{H}$
N99$^{H}$
G96$^{H}$
K94$^{H}$
V100a$^{H}$
Library 2 Size: **2x10$^{9}$**
C
3-pTza
H91$^{L}$
Y96$^{L}$
W28$^{L}$
T52$^{H}$
N32$^{L}$
Y50$^{H}$
S97$^{H}$
A53$^{H}$
G33$^{H}$
N99$^{H}$
L95$^{H}$
G96$^{H}$
Y32$^{H}$
V100a$^{H}$
K94$^{H}$
Library 3 Size: **4x10$^{9}$**
D
3-pTza
L95$^{H}$
S97$^{H}$
G98$^{H}$
N99$^{H}$
K94$^{H}$
G96$^{H}$
V100a$^{H}$
P100$^{H}$
Library 4 Size: **2x10$^{10}$**
E
N95a$^{L}$
E95$^{L}$
D95b$^{L}$
S94$^{L}$
A95c$^{L}$
G93$^{L}$
W28$^{L}$
R56$^{H}$
R29$^{L}$
T52$^{H}$
Y96$^{L}$
H91$^{L}$
3-pTza
Library 5 Size: **2x10$^{9}$**
F
3-pTza
F34$^{H}$
R71$^{H}$
Y50$^{H}$
V78$^{H}$
S35$^{H}$
F58$^{H}$
G49$^{H}$
I48$^{H}$
S67$^{H}$
H47$^{H}$
S61$^{H}$
A63$^{H}$
S65$^{H}$
W62$^{H}$
Library 6 Size: **3x10$^{9}$**

### Fig. S4.

**A**

| | | LFR1 | | | | | | | | | | | | | | | | | | | | | | |
|---|---|---|---|---|---|---|---|---|---|---|---|---|---|---|---|---|---|---|---|---|---|---|---|---|
| | KABAT | 1 | 2 | 3 | 4 | 5 | 6 | 7 | 8 | 9 | 10 | 11 | 12 | 13 | 14 | 15 | 16 | 17 | 18 | 19 | 20 | 21 | 22 | 23 |
| Scaffold | rSC44 | | Q | V | L | T | Q | T | P | S | F | T | S | A | A | V | G | G | T | V | T | I | N | C |
| | hSC44 | D | I | Q | | | | S | | | S | L | | | S | | | D | R | | | | T | |
| Library | hSC44 1 | | | | | | | | | | | | | | | | | | | | | | | |
| | hSC44 2 | | | | | | | | | | | | | | | | | | | | | | | |
| | hSC44 3 | | | | | | | | | | | | | | | | | | | | | | | |
| | hSC44 4 | | | | | | | | | | | | | | | | | | | | | | | |
| | hSC44 5 | | | | | | | | | | | | | | | | | | | | | | | |
| | hSC44 6 | | | | | | | | | | | | | | | | | | | | | | | |

| | | CDR L1 | | | | | | | | | | | | | LFR2 | | | | | | | | | | | | | | | CDR L2 | | | | | | |
|---|---|---|---|---|---|---|---|---|---|---|---|---|---|---|---|---|---|---|---|---|---|---|---|---|---|---|---|---|---|---|---|---|---|---|---|---|
| | KABAT | 24 | 25 | 26 | 27 | 27A | 27B | 28 | 29 | 30 | 31 | 32 | 33 | 34 | 35 | 36 | 37 | 38 | 39 | 40 | 41 | 42 | 43 | 44 | 45 | 46 | 47 | 48 | 49 | 50 | 51 | 52 | 53 | 54 | 55 | 56 |
| Scaffold | rSC44 | Q | S | S | Q | S | V | W | R | N | K | N | L | A | W | Y | Q | Q | K | P | G | Q | P | P | K | R | L | I | Y | A | I | A | T | L | D | S |
| | hSC44 | R | A | | | | | | | | | | V | | | | | | | | | K | A | | | | | | | | | | S | | Y | |
| Library | hSC44 1 | | | | | | | **W** | | | | **N** | | | | | | | | | | | | | | | | | | | | | | | | |
| | hSC44 2 | | | | | | | **W** | **R** | | **K** | **N** | | | | | | | | | | | | | | | | | | | | | | | | |
| | hSC44 3 | | | | | | | **W** | | | | **N** | | | | | | | | | | | | | | | | | | | | | | | | |
| | hSC44 4 | | | | | | | | | | | | | | | | | | | | | | | | | | | | | | | | | | | |
| | hSC44 5 | | | | | | | **W** | **R** | | | | | | | | | | | | | | | | | | | | | | | | | | | |
| | hSC44 6 | | | | | | | | | | | | | | | | | | | | | | | | | | | | | | | | | | | |

| | | LFR3 | | | | | | | | | | | | | | | | | | | | | | | | | | | | | | |
|---|---|---|---|---|---|---|---|---|---|---|---|---|---|---|---|---|---|---|---|---|---|---|---|---|---|---|---|---|---|---|---|---|---|
| | KABAT | 57 | 58 | 59 | 60 | 61 | 62 | 63 | 64 | 65 | 66 | 67 | 68 | 69 | 70 | 71 | 72 | 73 | 74 | 75 | 76 | 77 | 78 | 79 | 80 | 81 | 82 | 83 | 84 | 85 | 86 | 87 | 88 |
| Scaffold | rSC44 | G | V | P | S | R | F | S | G | S | G | S | G | T | Q | F | T | L | T | I | S | D | V | Q | C | D | D | A | A | T | Y | Y | C |
| | hSC44 | | | | | | | | | | | | | | D | | | | | | | S | L | | P | E | | F | | | | | |
| Library | hSC44 1 | | | | | | | | | | | | | | | | | | | | | | | | | | | | | | | | |
| | hSC44 2 | | | | | | | | | | | | | | | | | | | | | | | | | | | | | | | | |
| | hSC44 3 | | | | | | | | | | | | | | | | | | | | | | | | | | | | | | | | |
| | hSC44 4 | | | | | | | | | | | | | | | | | | | | | | | | | | | | | | | | |
| | hSC44 5 | | | | | | | | | | | | | | | | | | | | | | | | | | | | | | | | |
| | hSC44 6 | | | | | | | | | | | | | | | | | | | | | | | | | | | | | | | | |

| | | CDR L3 | | | | | | | | | | | | | LFR4 | | | | | | | | | |
|---|---|---|---|---|---|---|---|---|---|---|---|---|---|---|---|---|---|---|---|---|---|---|---|---|
| | KABAT | 89 | 90 | 91 | 92 | 93 | 94 | 95 | 95a | 95b | 95c | 95d | 96 | 97 | 98 | 99 | 100 | 101 | 102 | 103 | 104 | 105 | 106 | 107 |
| Scaffold | rSC44 | V | G | H | Y | G | S | E | N | D | A | Y | Y | A | F | G | G | G | T | E | V | V | V | K |
| | hSC44 | | | | | | | | | | | | | | | | Q | | | K | | E | I | |
| Library | hSC44 1 | | | **H** | | | | | | | | | **Y** | | | | | | | | | | | |
| | hSC44 2 | | | **H** | | | | | | | | | **Y** | | | | | | | | | | | |
| | hSC44 3 | | | **H** | | | | | | | | | **Y** | | | | | | | | | | | |
| | hSC44 4 | | | | | | | | | | | | | | | | | | | | | | | |
| | hSC44 5 | | | **H** | | **G** | **S** | **E** | **N** | **D** | **A** | | **Y** | | | | | | | | | | | |
| | hSC44 6 | | | | | | | | | | | | | | | | | | | | | | | |

**B**

| | | HFR1 | | | | | | | | | | | | | | | | | | | | | | | | | | | | |
|---|---|---|---|---|---|---|---|---|---|---|---|---|---|---|---|---|---|---|---|---|---|---|---|---|---|---|---|---|---|---|---|
| | KABAT | 1 | 2 | 3 | 4 | 5 | 6 | 7 | 8 | 9 | 10 | 11 | 12 | 13 | 14 | 15 | 16 | 17 | 18 | 19 | 20 | 21 | 22 | 23 | 24 | 25 | 26 | 27 | 28 | 29 | 30 |
| Scaffold | rSC44 | | Q | S | V | K | E | S | G | R | G | L | V | Q | P | G | G | S | L | T | L | T | C | T | A | S | G | F | S | I | D |
| | hSC44 | E | V | Q | L | V | | | | G | | | | | | | | | | R | | S | | | | | | | | | |
| Library | hSC44 1 | | | | | | | | | | | | | | | | | | | | | | | | | | | | | | |
| | hSC44 2 | | | | | | | | | | | | | | | | | | | | | | | | | | | | | | |
| | hSC44 3 | | | | | | | | | | | | | | | | | | | | | | | | | | | | | | |
| | hSC44 4 | | | | | | | | | | | | | | | | | | | | | | | | | | | | | | |
| | hSC44 5 | | | | | | | | | | | | | | | | | | | | | | | | | | | | | | |
| | hSC44 6 | | | | | | | | | | | | | | | | | | | | | | | | | | | | | | |

| | | CDR H1 | | | | | HFR2 | | | | | | | | | | | | | | CDR H2 | | | | | | | | | | | | | | | |
|---|---|---|---|---|---|---|---|---|---|---|---|---|---|---|---|---|---|---|---|---|---|---|---|---|---|---|---|---|---|---|---|---|---|---|---|---|
| | KABAT | 31 | 32 | 33 | 34 | 35 | 36 | 37 | 38 | 39 | 40 | 41 | 42 | 43 | 44 | 45 | 46 | 47 | 48 | 49 | 50 | 51 | 52 | 53 | 54 | 55 | 56 | 57 | 58 | 59 | 60 | 61 | 62 | 63 | 64 | 65 |
| Scaffold | rSC44 | S | Y | G | F | S | W | V | R | Q | A | P | G | K | G | L | E | H | I | G | Y | L | T | A | G | G | R | A | F | Y | A | S | W | A | K | S |
| | hSC44 | | | | | | | | | | | | | | | | | | | | | | | | | | | | | | | | | | | |
| Library | hSC44 1 | | | | | | | | | | | | | | | | | | | | | | | | | | | | | | | | | | | |
| | hSC44 2 | | **Y** | **G** | | | | | | | | | | | | | | | | | | | | | | | | | | | | | | | | |
| | hSC44 3 | | | | | | | | | | | | | | | | | | | | | | **T** | **A** | | | | | | | | | | | | |
| | hSC44 4 | | **Y** | **G** | | | | | | | | | | | | | | | | | | | | | | | | | | | | | | | | |
| | hSC44 5 | | | | | | | | | | | | | | | | | | | | | | **T** | | | | **R** | | | | | | | | | |
| | hSC44 6 | | | | **F** | **S** | | | | | | | | | | | | **H** | **I** | **G** | **Y** | | | | | | | | **F** | | | **S** | **W** | **A** | | **S** |

| | | HFR3 | | | | | | | | | | | | | | | | | | | | | | | | | | | | | | |
|---|---|---|---|---|---|---|---|---|---|---|---|---|---|---|---|---|---|---|---|---|---|---|---|---|---|---|---|---|---|---|---|---|
| | KABAT | 66 | 67 | 68 | 69 | 70 | 71 | 72 | 73 | 74 | 75 | 76 | 77 | 78 | 79 | 80 | 81 | 82 | 82A | 82B | 82C | 83 | 84 | 85 | 86 | 87 | 88 | 89 | 90 | 91 | 92 | 93 | 94 |
| Scaffold | rSC44 | R | S | T | I | T | R | N | T | N | E | N | T | V | T | L | K | M | T | S | L | T | A | A | D | T | A | T | Y | F | C | A | K |
| | hSC44 | | | | | | | | | | | | | | | | | | N | | | R | | E | | | | V | | Y | | | |
| Library | hSC44 1 | | | | | | | | | | | | | | | | | | | | | | | | | | | | | | | | **K** |
| | hSC44 2 | | | | | | | | | | | | | | | | | | | | | | | | | | | | | | | | **K** |
| | hSC44 3 | | | | | | | | | | | | | | | | | | | | | | | | | | | | | | | | **K** |
| | hSC44 4 | | | | | | | | | | | | | | | | | | | | | | | | | | | | | | | | **K** |
| | hSC44 5 | | | | | | | | | | | | | | | | | | | | | | | | | | | | | | | | |
| | hSC44 6 | | **S** | | | | **R** | | | | | | | **V** | | | | | | | | | | | | | | | | | | | |

| | | CDR H3 | | | | | | | | | HFR4 | | | | | | | | | | |
|---|---|---|---|---|---|---|---|---|---|---|---|---|---|---|---|---|---|---|---|---|---|
| | KABAT | 95 | 96 | 97 | 98 | 99 | 100 | 100a | 101 | 102 | 103 | 104 | 105 | 106 | 107 | 108 | 109 | 110 | 111 | 112 | 113 |
| Scaffold | rSC44 | L | G | S | G | N | P | V | A | I | W | G | P | G | T | L | V | T | V | S | S |
| | hSC44 | | | | | | | | | | | | Q | | | | | | V | S | S |
| Library | hSC44 1 | **L** | **G** | **S** | | **N** | | **V** | | | | | | | | | | | | | |
| | hSC44 2 | **L** | **G** | **S** | | **N** | | **V** | | | | | | | | | | | | | |
| | hSC44 3 | **L** | **G** | **S** | | **N** | | **V** | | | | | | | | | | | | | |
| | hSC44 4 | **L** | **G** | **S** | **G** | **N** | **P** | **V** | | | | | | | | | | | | | |
| | hSC44 5 | | | | | | | | | | | | | | | | | | | | |
| | hSC44 6 | | | | | | | | | | | | | | | | | | | | |

**Fig. S5.**

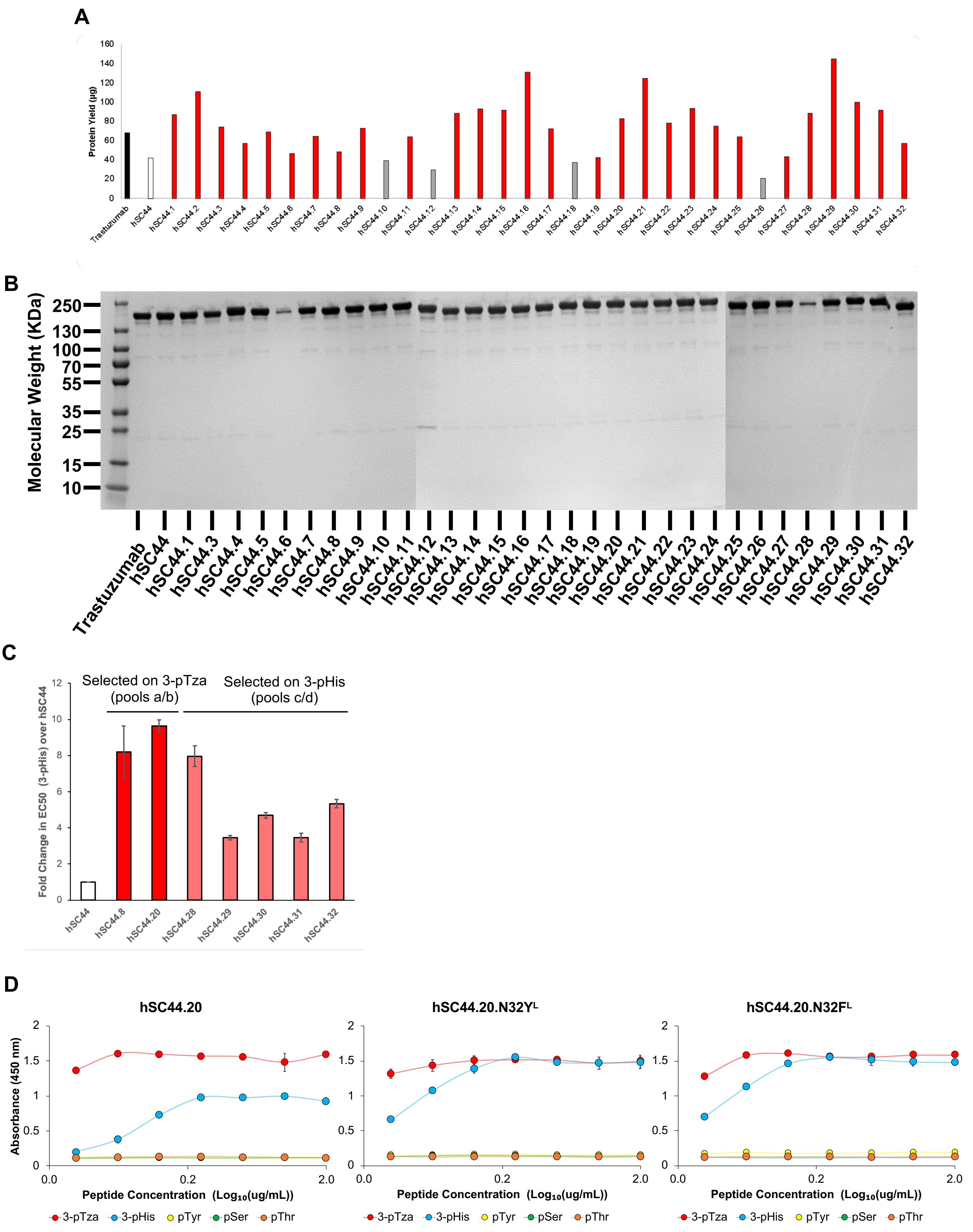
A
Protein Yield (µg)
Trastuzumab
hSC44
hSC44.1
hSC44.2
hSC44.3
hSC44.4
hSC44.5
hSC44.6
hSC44.7
hSC44.8
hSC44.9
hSC44.10
hSC44.11
hSC44.12
hSC44.13
hSC44.14
hSC44.15
hSC44.16
hSC44.17
hSC44.18
hSC44.19
hSC44.20
hSC44.21
hSC44.22
hSC44.23
hSC44.24
hSC44.25
hSC44.26
hSC44.27
hSC44.28
hSC44.29
hSC44.30
hSC44.31
hSC44.32
B
Molecular Weight (KDa)
250
130
100
70
55
35
25
15
10
C
Selected on 3-pTza (pools a/b)
Selected on 3-pHis (pools c/d)
Fold Change in EC50 (3-pHis) over hSC44
D
hSC44.20
hSC44.20.N32Y^L
hSC44.20.N32F^L
Absorbance (450 nm)
Peptide Concentration (Log10(ug/mL))
3-pTza
3-pHis
pTyr
pSer
pThr

**Fig. S6.**

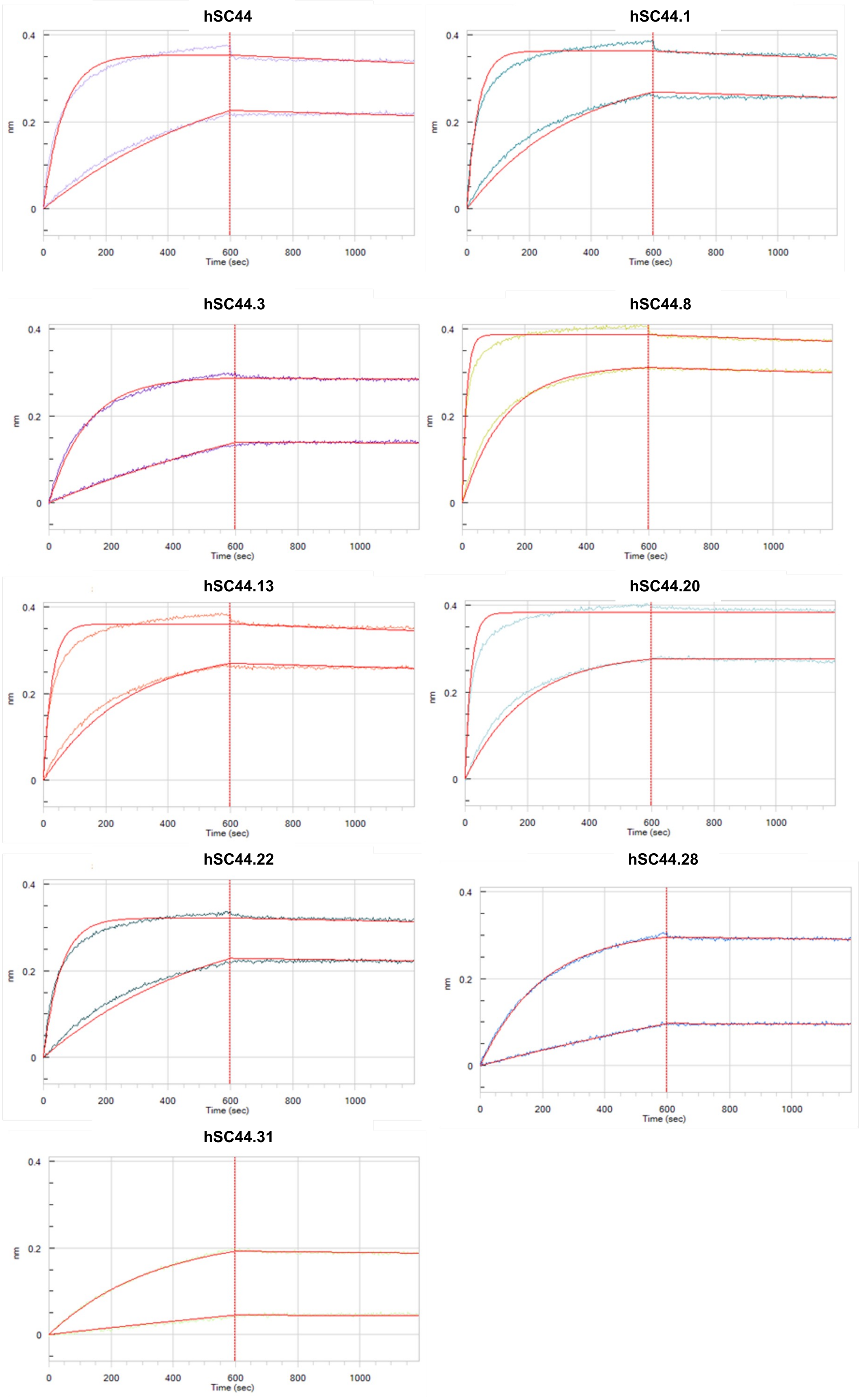
hSC44
hSC44.1
hSC44.3
hSC44.8
hSC44.13
hSC44.20
hSC44.22
hSC44.28
hSC44.31
nm
0.4
0.2
0
Time (sec)
0
200
400
600
800
1000

**Fig. S7.**

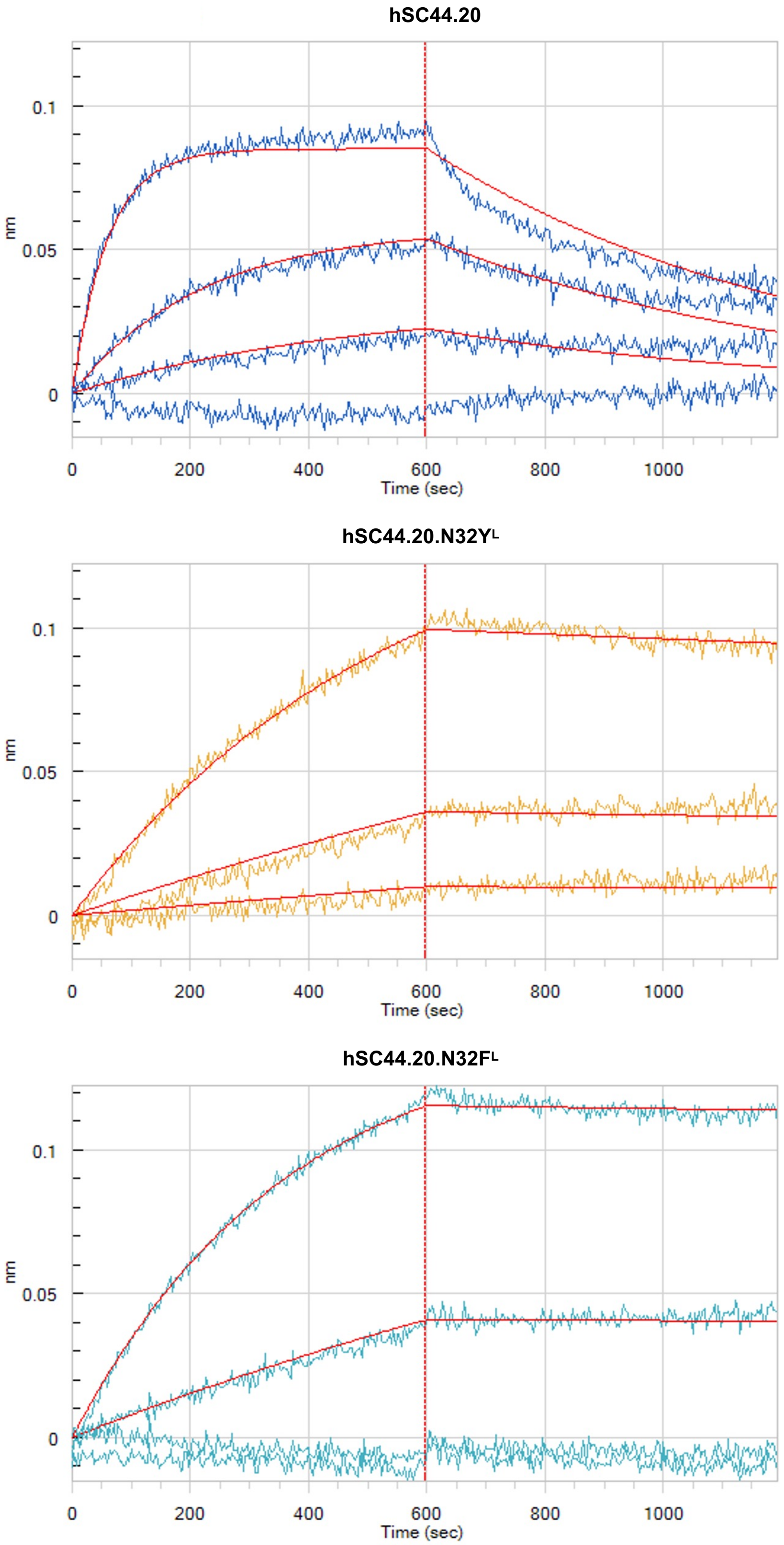
hSC44.20
0.1
0.05
0
nm
0
200
400
600
800
1000
Time (sec)
hSC44.20.N32Y^L
0.1
0.05
0
nm
0
200
400
600
800
1000
Time (sec)
hSC44.20.N32F^L
0.1
0.05
0
nm
0
200
400
600
800
1000
Time (sec)

**Fig. S8.**

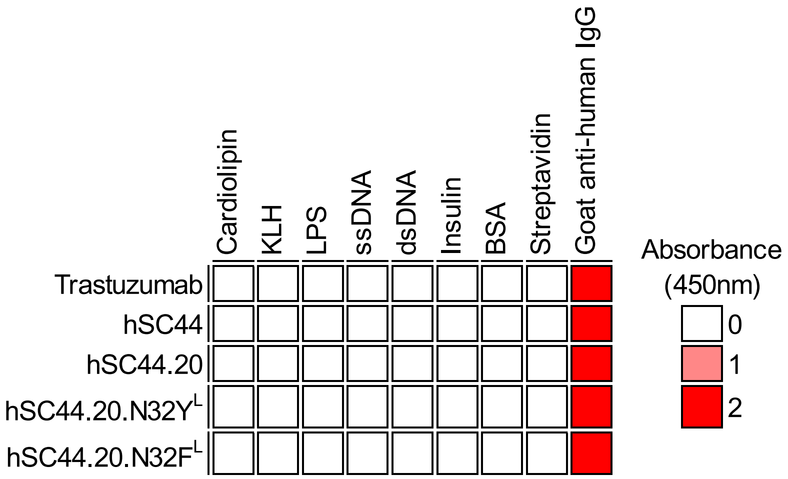
Cardiolipin
KLH
LPS
ssDNA
dsDNA
Insulin
BSA
Streptavidin
Goat anti-human IgG
Trastuzumab
hSC44
hSC44.20
hSC44.20.N32Y^L
hSC44.20.N32F^L
Absorbance
(450nm)
0
1
2

**Fig. S9.**

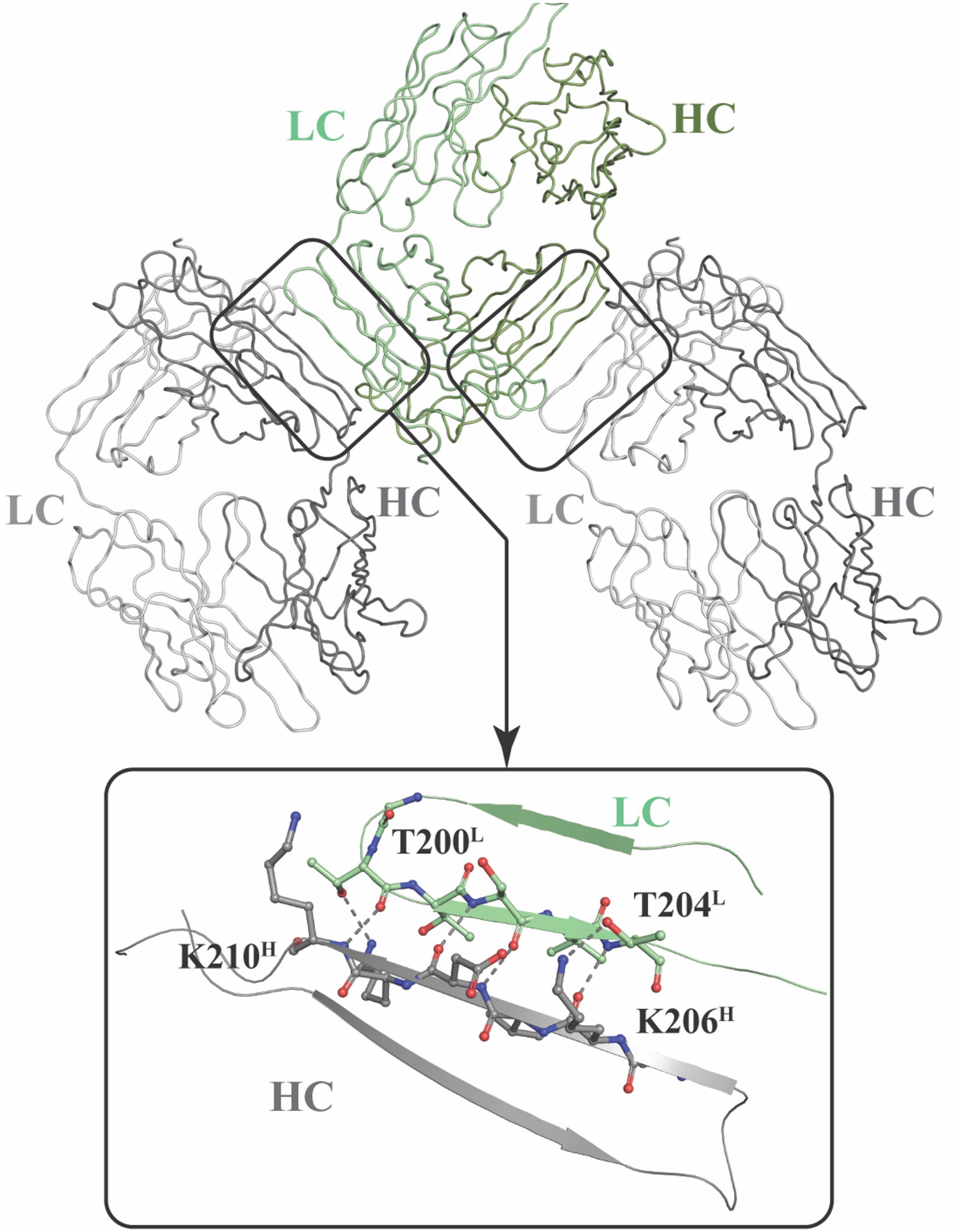

**Fig. S10.**

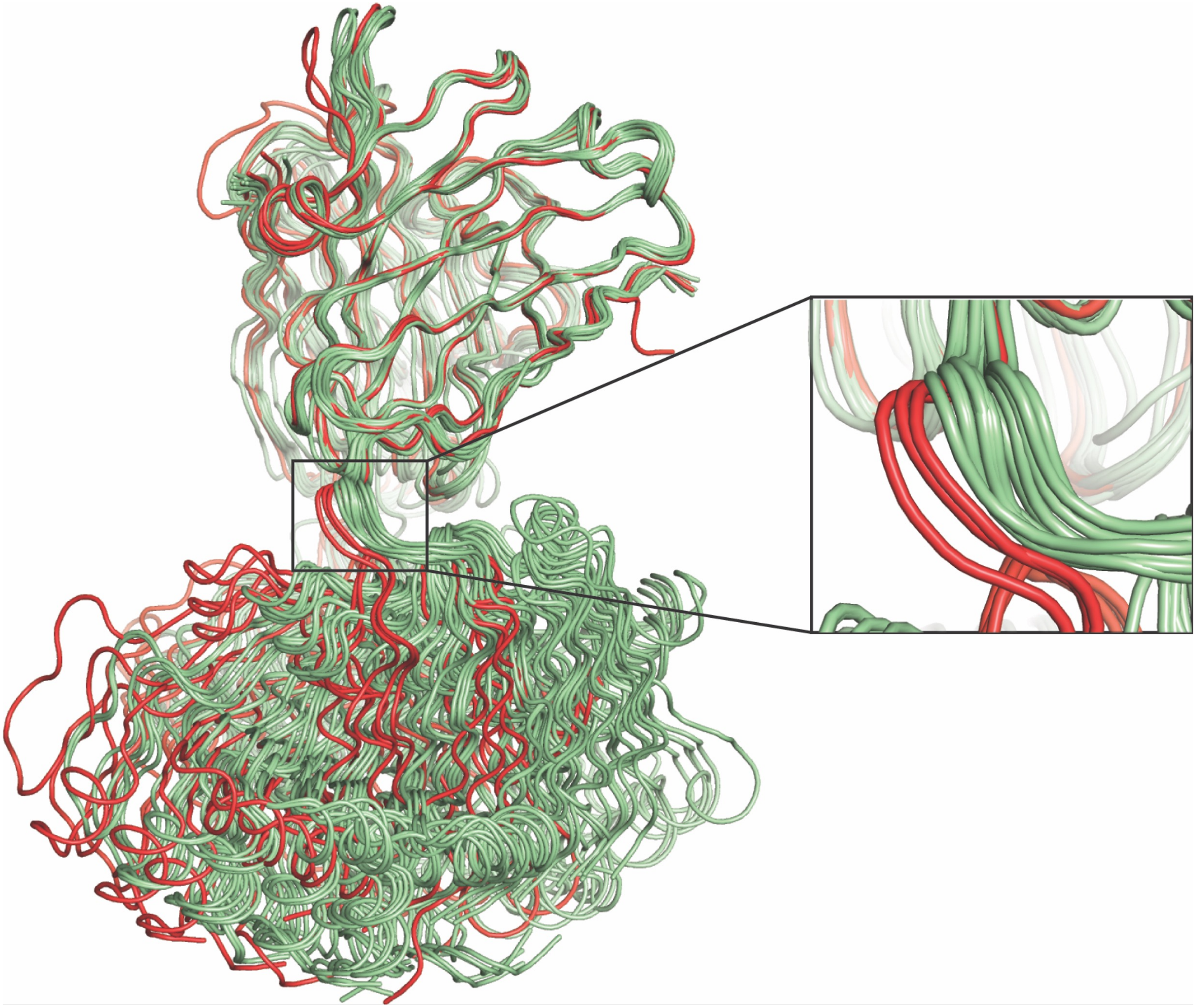

**Fig. S11.**

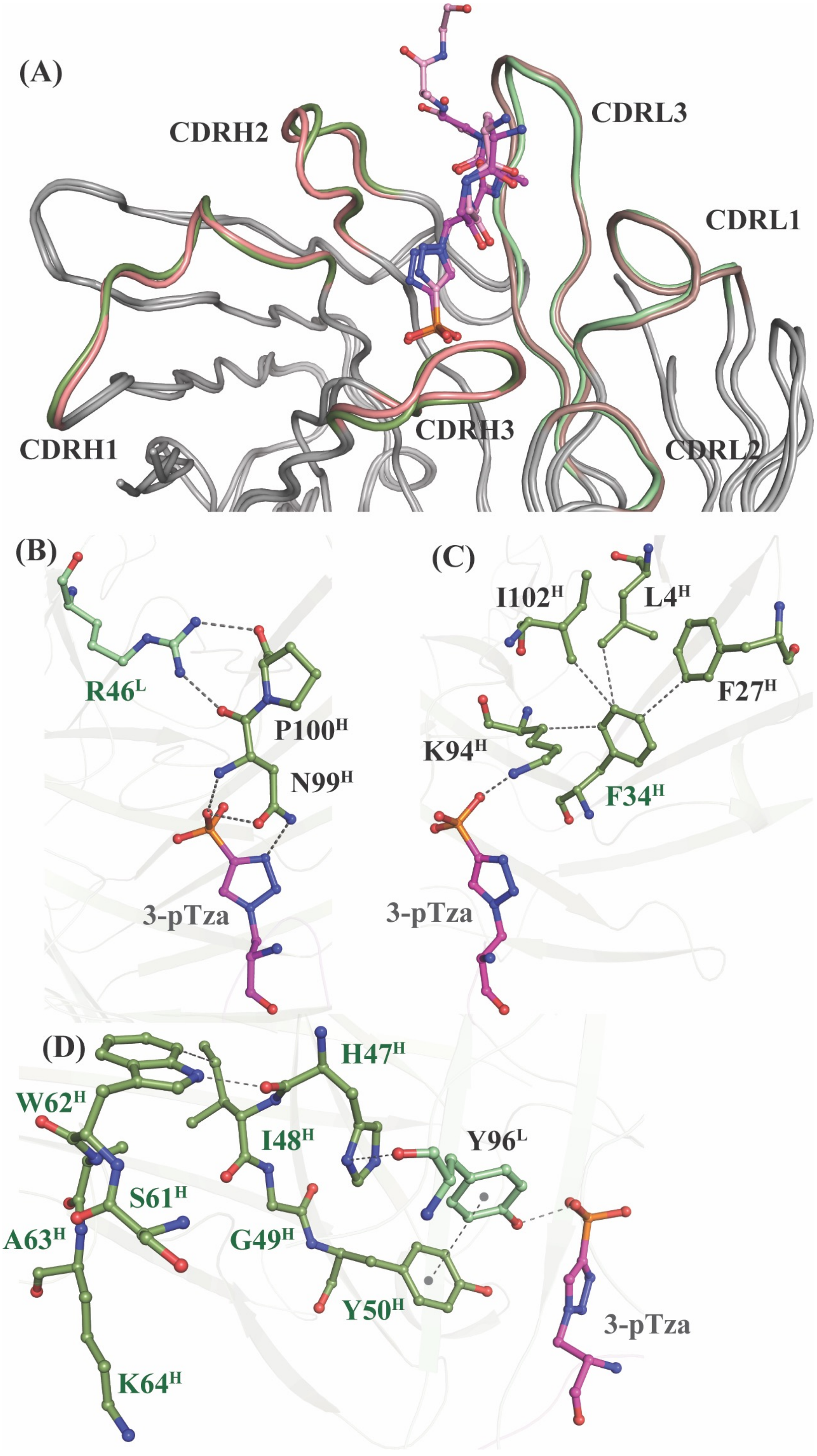
(A)
CDRH2
CDRL3
CDRL1
CDRH1
CDRH3
CDRL2
(B)
R46L
P100H
N99H
3-pTza
(C)
I102H
L4H
F27H
K94H
F34H
3-pTza
(D)
H47H
W62H
I48H
Y96L
S61H
A63H
G49H
Y50H
3-pTza
K64H

**Fig. S12.**

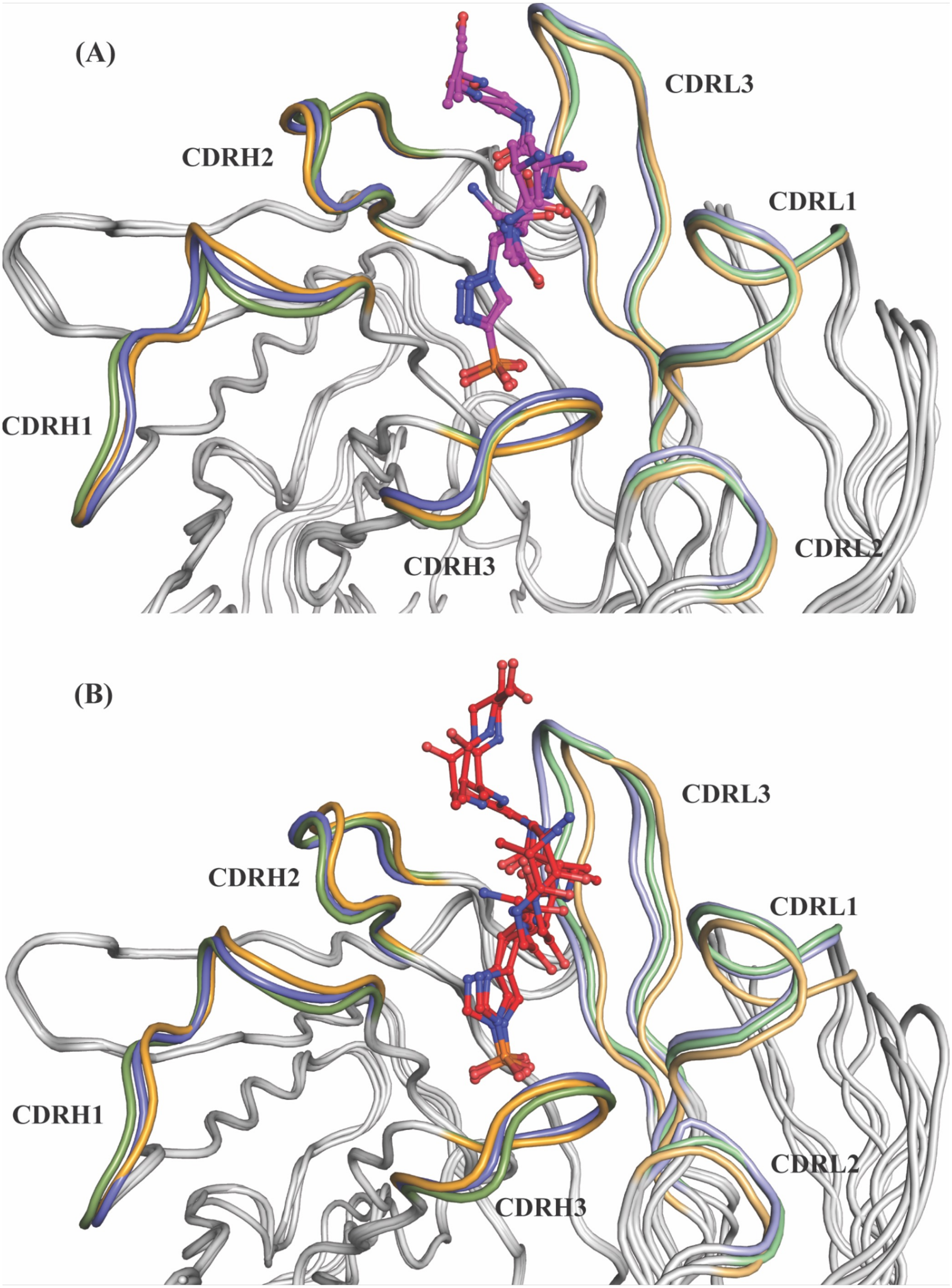
(A)
CDRL3
CDRH2
CDRL1
CDRH1
CDRH3
CDRL2
(B)
CDRL3
CDRH2
CDRL1
CDRH1
CDRL2
CDRH3

**Fig. S13.**

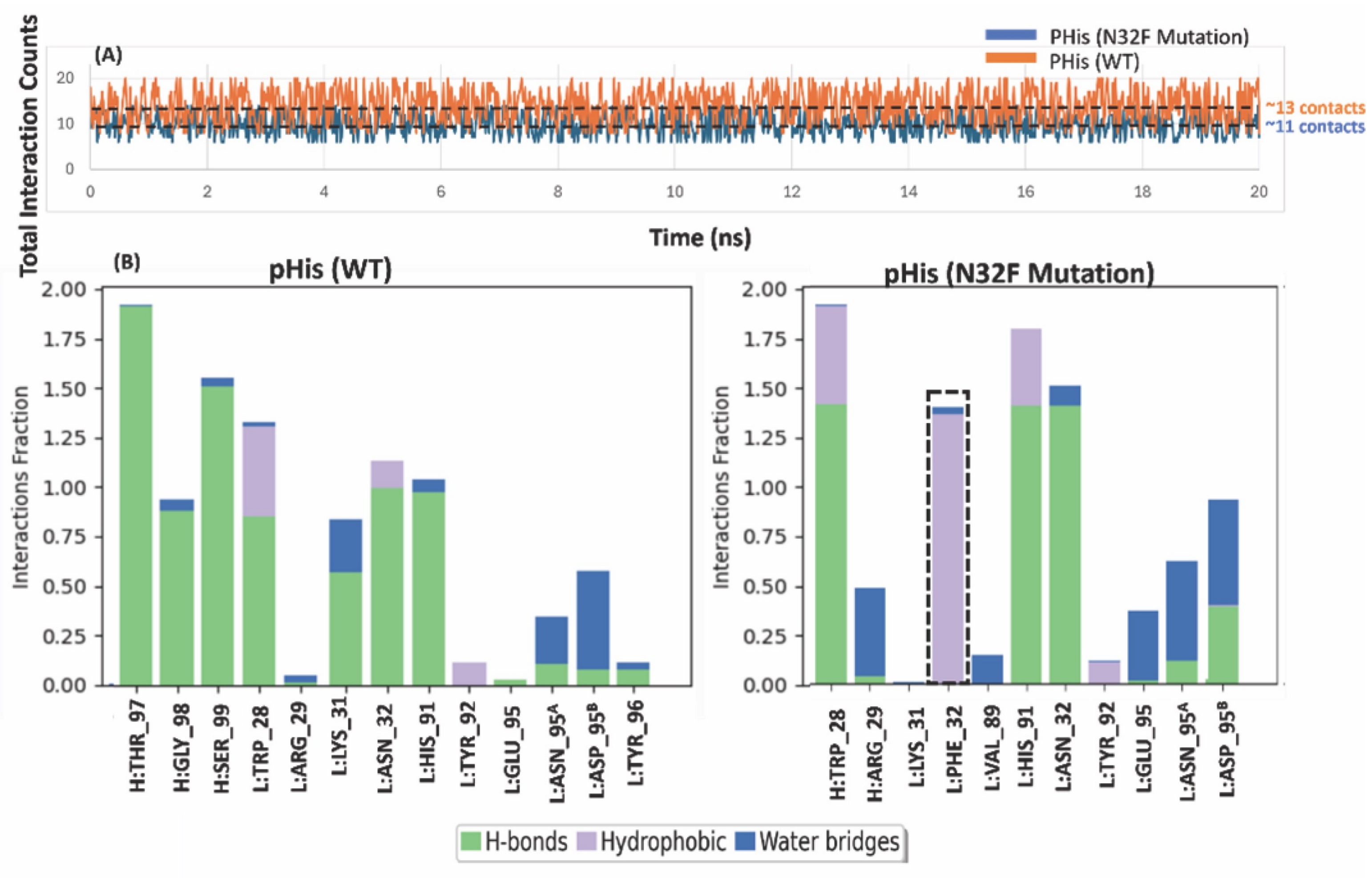

(A)
PHis (N32F Mutation)
PHis (WT)
~13 contacts
~11 contacts
Total Interaction Counts
Time (ns)
(B)
pHis (WT)
pHis (N32F Mutation)
Interactions Fraction
H:THR_97
H:GLY_98
H:SER_99
L:TRP_28
L:ARG_29
L:LYS_31
L:ASN_32
L:HIS_91
L:TYR_92
L:GLU_95
L:ASN_95A
L:ASP_95B
L:TYR_96
H:TRP_28
H:ARG_29
L:PHE_32
L:VAL_89
H-bonds
Hydrophobic
Water bridges

**Fig. S14.**

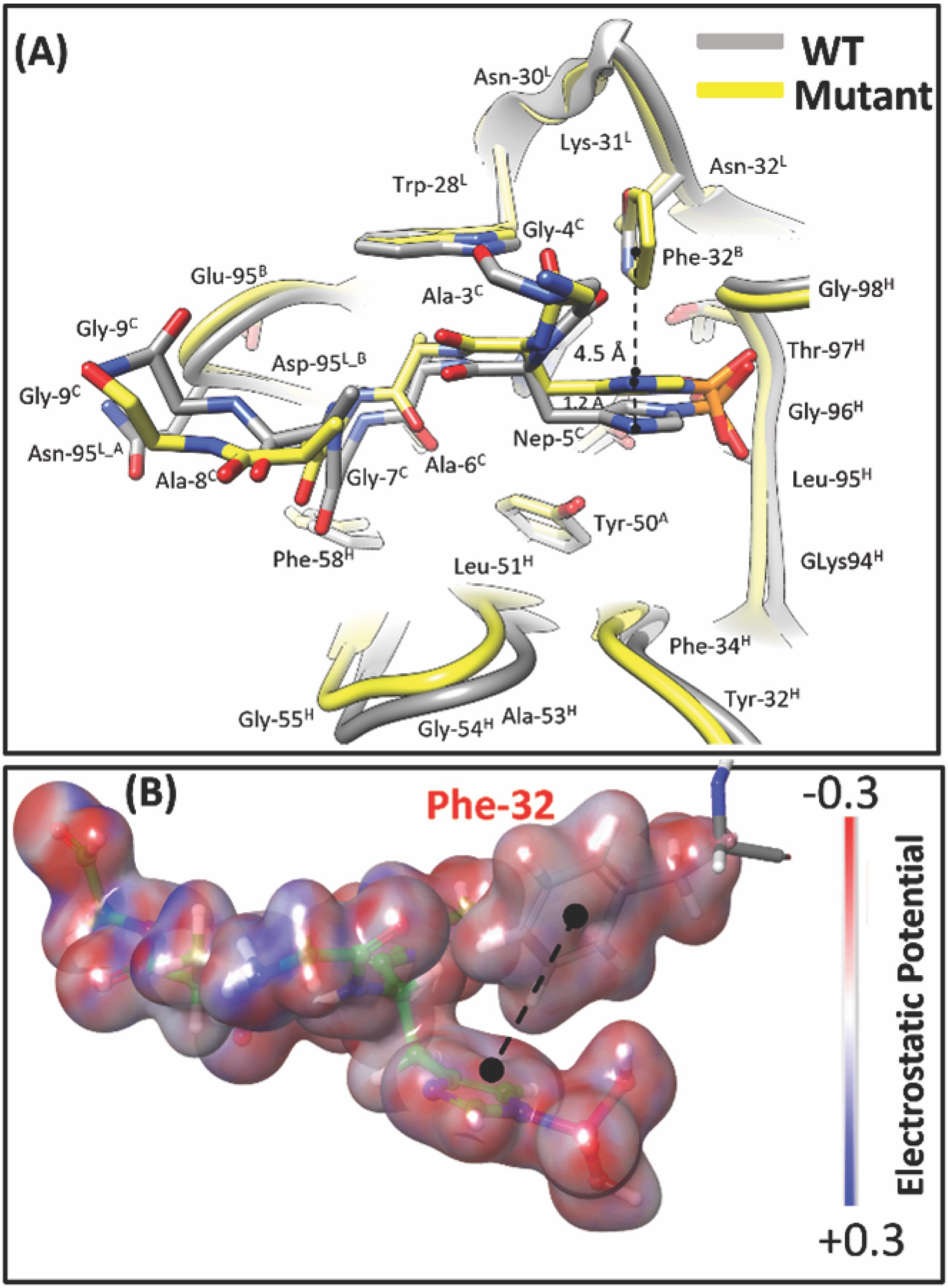
(A)
WT
Mutant
Asn-30L
Lys-31L
Asn-32L
Trp-28L
Gly-4C
Phe-32B
Glu-95B
Ala-3C
Gly-98H
Gly-9C
Asp-95L_B
4.5 Å
Thr-97H
Gly-9C
1.2 Å
Gly-96H
Nep-5C
Asn-95L_A
Ala-8C
Gly-7C
Ala-6C
Leu-95H
Tyr-50A
Phe-58H
Leu-51H
GLys94H
Phe-34H
Tyr-32H
Gly-55H
Gly-54H
Ala-53H
(B)
Phe-32
-0.3
Electrostatic Potential
+0.3

**Fig. S15.**

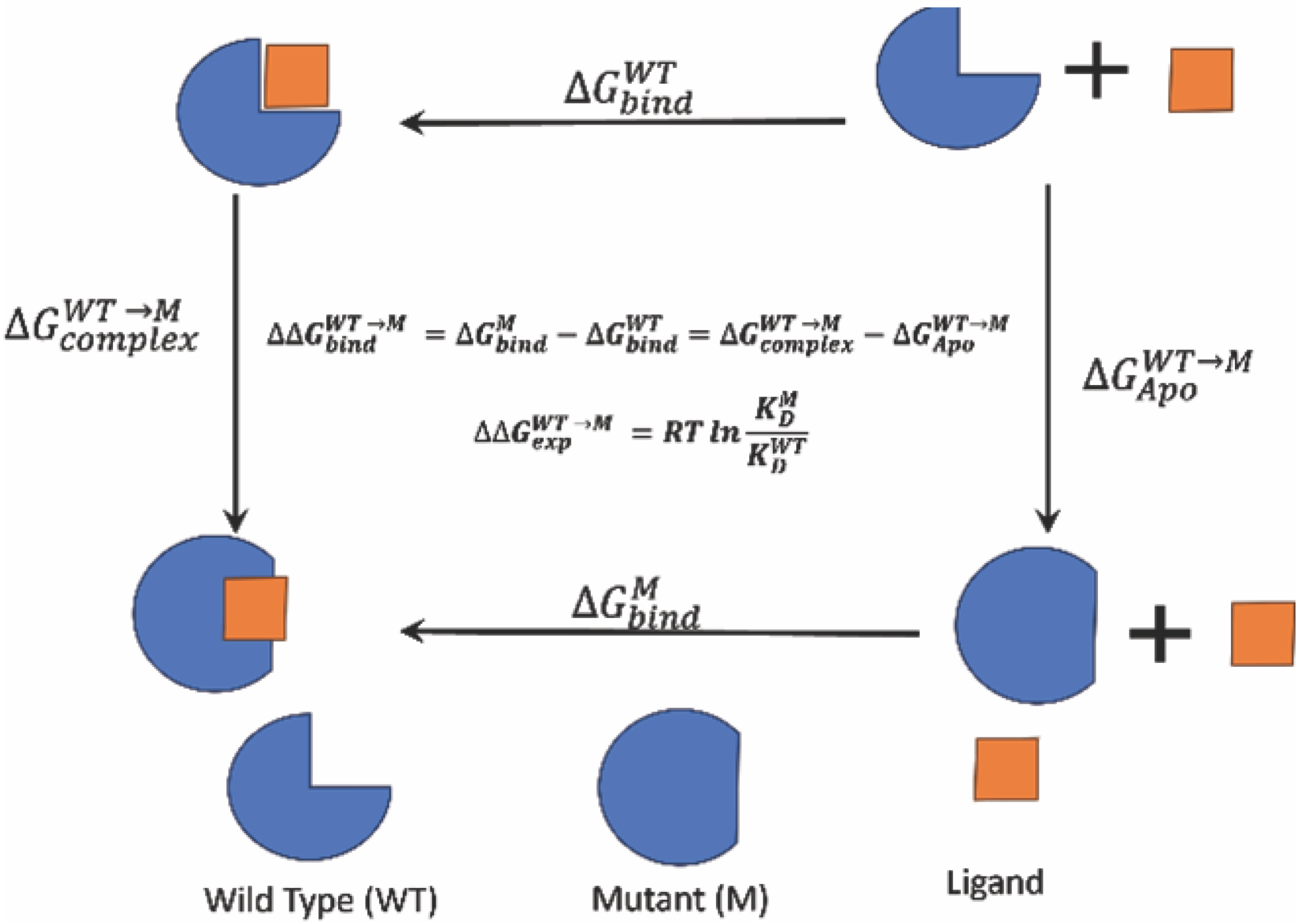

$\Delta G_{bind}^{WT}$
$\Delta G_{complex}^{WT\rightarrow M}$
$\Delta\Delta G_{bind}^{WT\rightarrow M} = \Delta G_{bind}^{M} - \Delta G_{bind}^{WT} = \Delta G_{complex}^{WT\rightarrow M} - \Delta G_{Apo}^{WT\rightarrow M}$
$\Delta\Delta G_{exp}^{WT\rightarrow M} = RT\, ln \frac{K_D^M}{K_D^{WT}}$
$\Delta G_{Apo}^{WT\rightarrow M}$
$\Delta G_{bind}^{M}$
+
+
Wild Type (WT)
Mutant (M)
Ligand

**Fig. S16.**

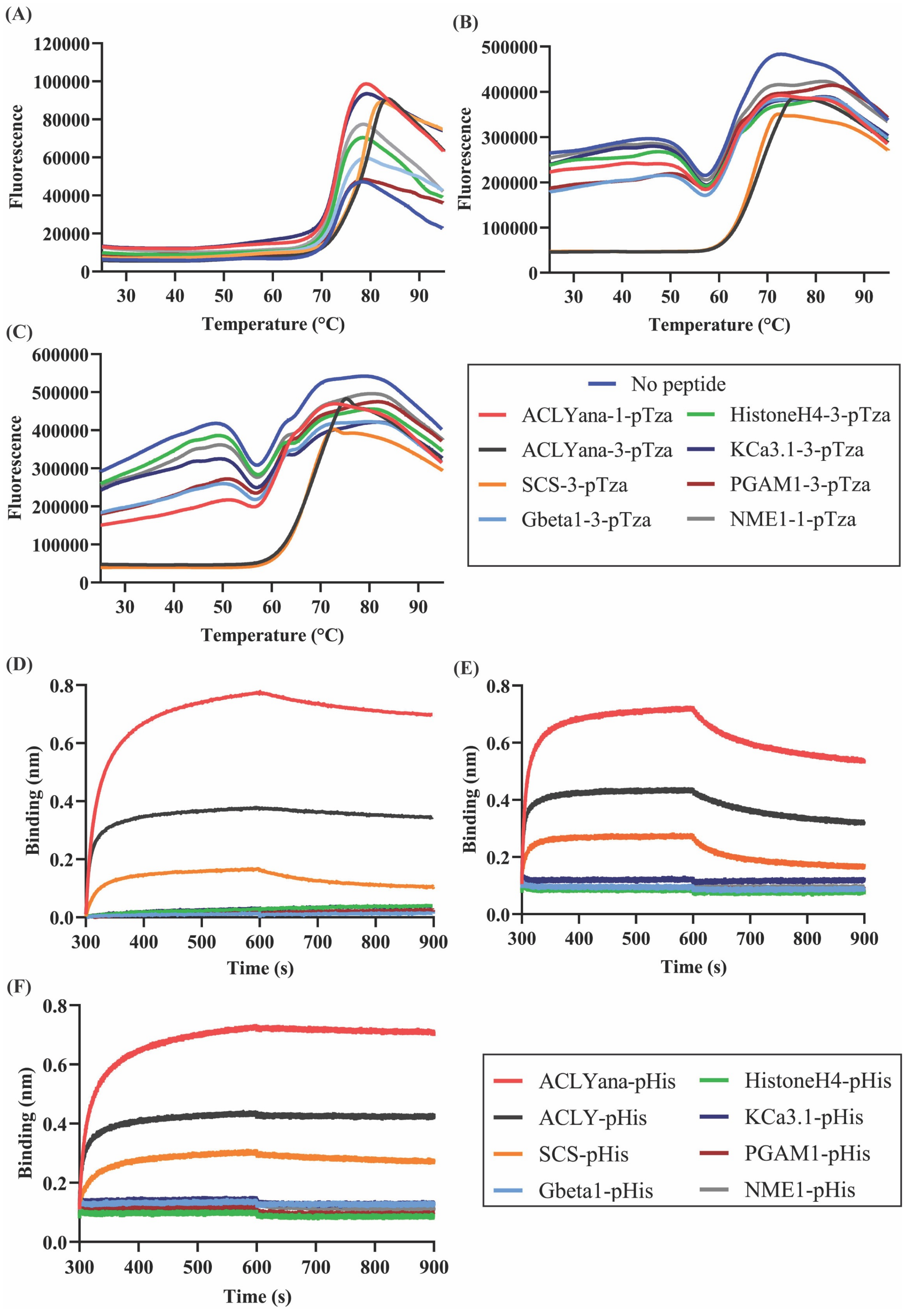
(A)
Fluorescence
Temperature (°C)
(B)
Fluorescence
Temperature (°C)
(C)
Fluorescence
Temperature (°C)
No peptide
ACLYana-1-pTza
ACLYana-3-pTza
SCS-3-pTza
Gbeta1-3-pTza
HistoneH4-3-pTza
KCa3.1-3-pTza
PGAM1-3-pTza
NME1-1-pTza
(D)
Binding (nm)
Time (s)
(E)
Binding (nm)
Time (s)
(F)
Binding (nm)
Time (s)
ACLYana-pHis
ACLY-pHis
SCS-pHis
Gbeta1-pHis
HistoneH4-pHis
KCa3.1-pHis
PGAM1-pHis
NME1-pHis

**Fig. S17.**

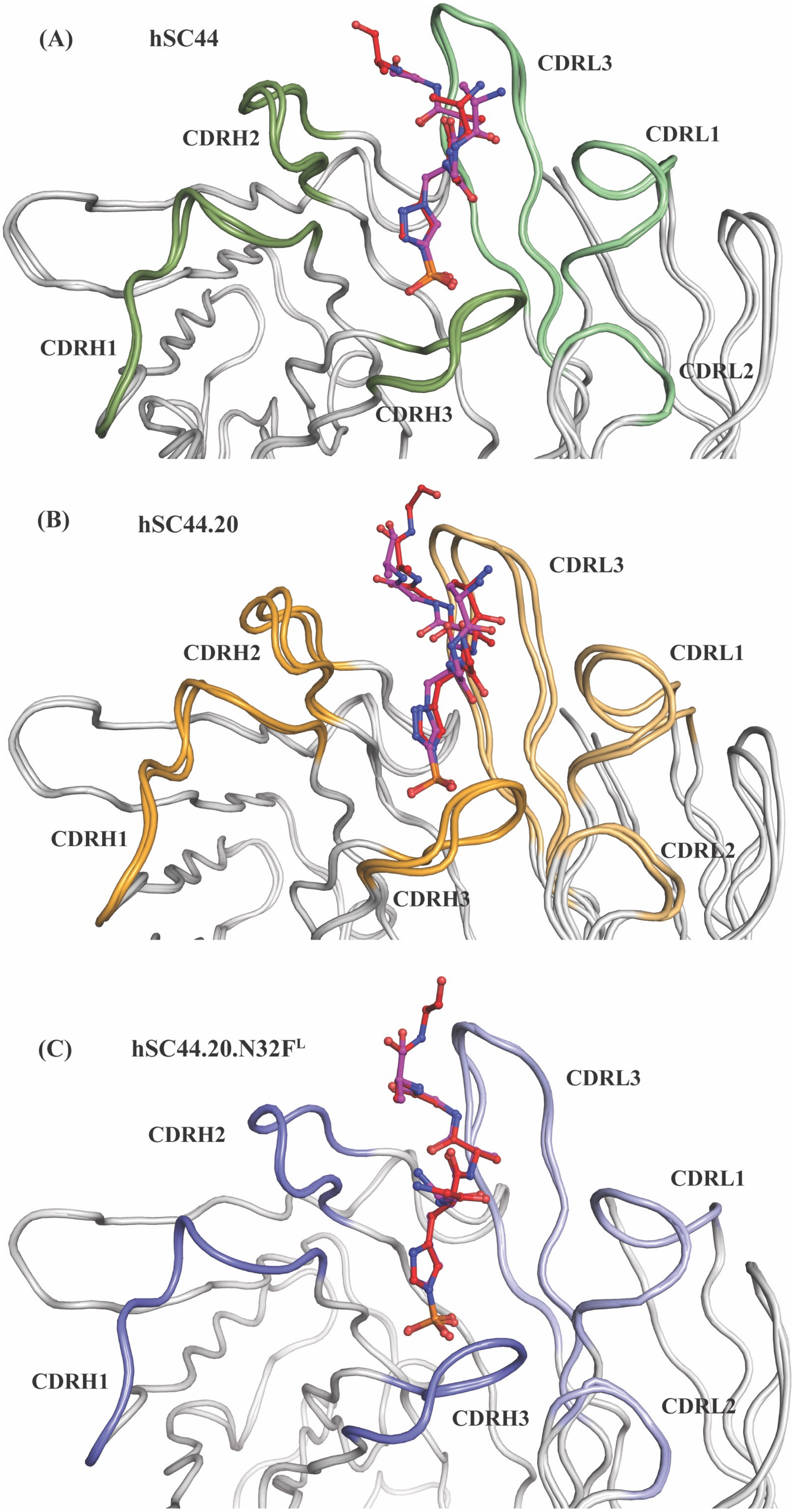
(A)
hSC44
CDRL3
CDRH2
CDRL1
CDRH1
CDRL2
CDRH3
(B)
hSC44.20
CDRL3
CDRH2
CDRL1
CDRH1
CDRL2
CDRH3
(C)
hSC44.20.N32F^L
CDRL3
CDRH2
CDRL1
CDRH1
CDRL2
CDRH3

**Fig. S18.**

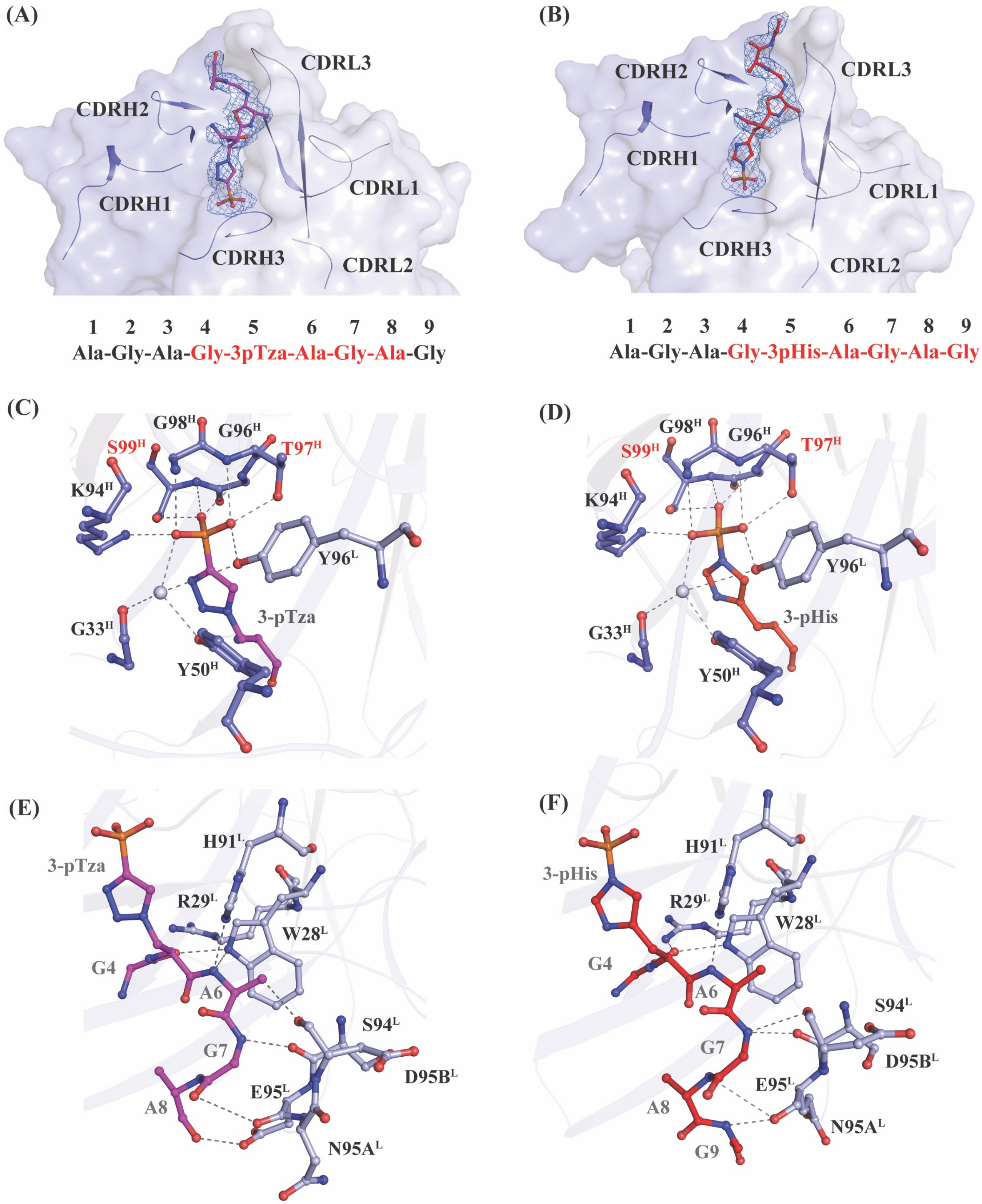

(A)
CDRL3
CDRH2
CDRH1
CDRL1
CDRH3
CDRL2
1 2 3 4 5 6 7 8 9
Ala-Gly-Ala-Gly-3pTza-Ala-Gly-Ala-Gly
(B)
CDRH2
CDRL3
CDRH1
CDRL1
CDRH3
CDRL2
1 2 3 4 5 6 7 8 9
Ala-Gly-Ala-Gly-3pHis-Ala-Gly-Ala-Gly
(C)
G98H
G96H
S99H
T97H
K94H
Y96L
3-pTza
G33H
Y50H
(D)
G98H
G96H
S99H
T97H
K94H
Y96L
3-pHis
G33H
Y50H
(E)
3-pTza
H91L
R29L
W28L
G4
A6
S94L
G7
D95BL
E95L
A8
N95AL
(F)
3-pHis
H91L
R29L
W28L
G4
A6
S94L
G7
D95BL
E95L
A8
N95AL
G9

**Fig. S19.**

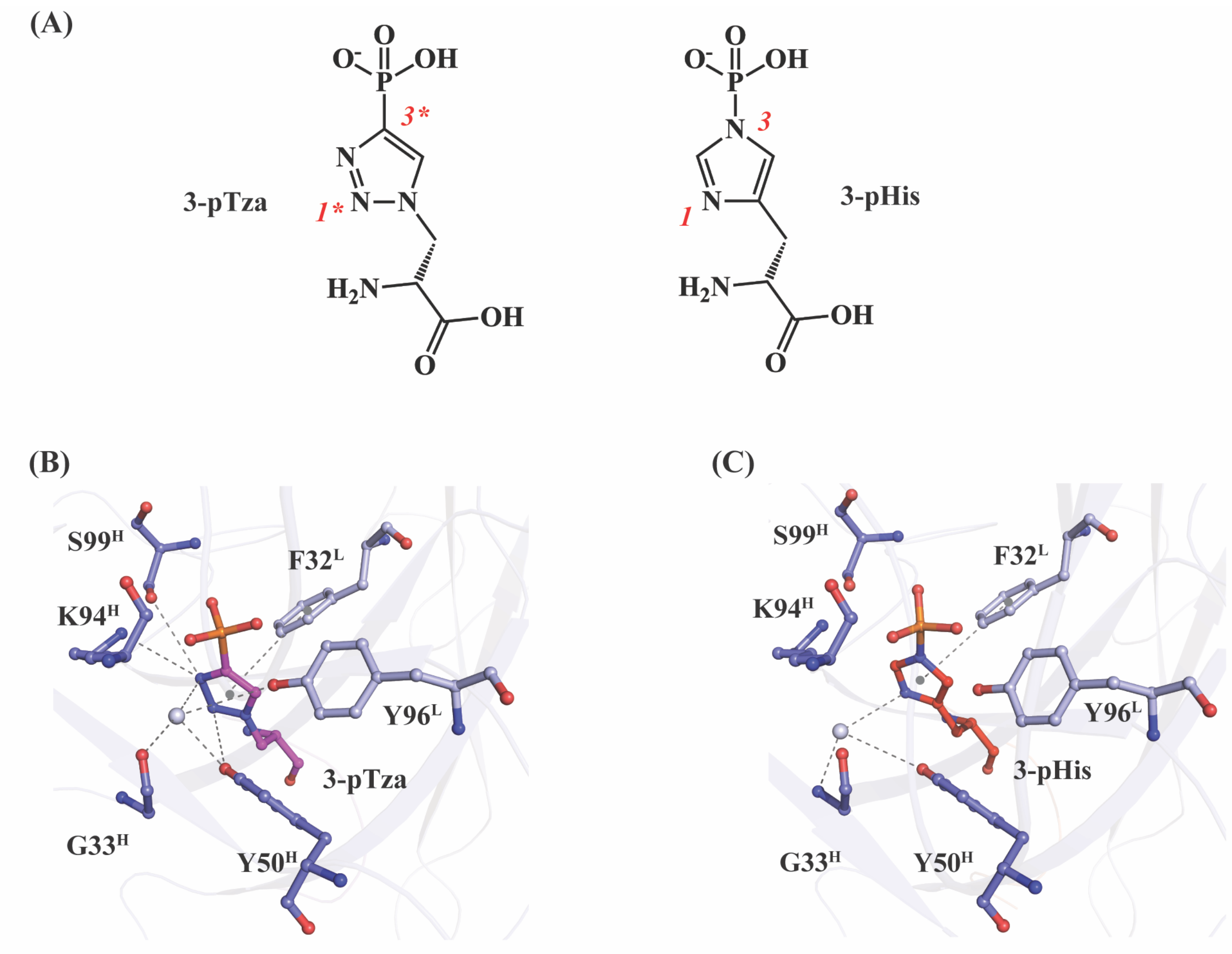

(A)
3-pTza
3*
1*
3-pHis
3
1
(B)
S99H
F32L
K94H
Y96L
3-pTza
G33H
Y50H
(C)
S99H
F32L
K94H
Y96L
3-pHis
G33H
Y50H

**Fig. S20.**

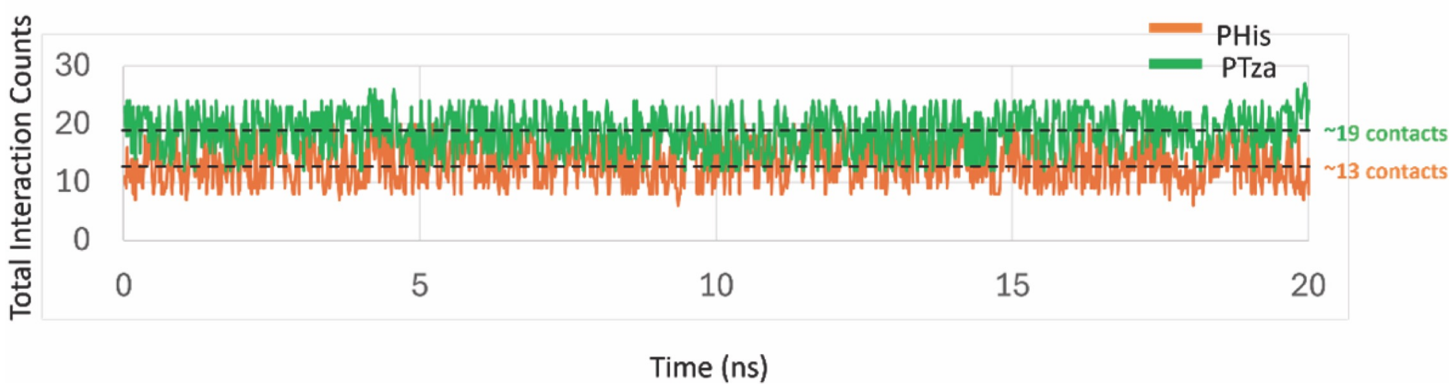

PHis
PTza
~19 contacts
~13 contacts
Total Interaction Counts
30
20
10
0
0
5
10
15
20
Time (ns)

**Fig. S21**

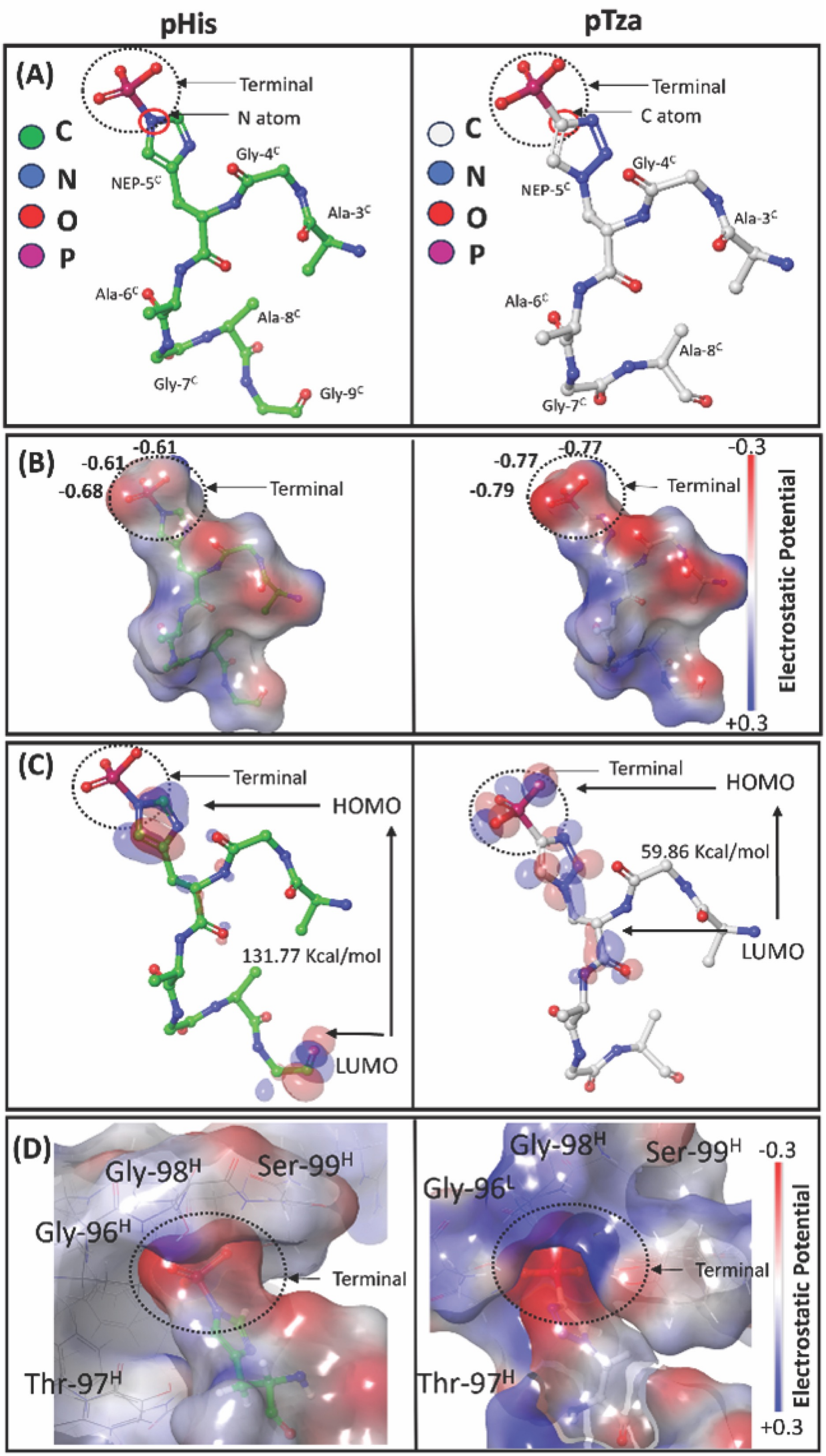
pHis
pTza
(A)
Terminal
N atom
C atom
C
N
O
P
NEP-5C
Gly-4C
Ala-3C
Ala-6C
Ala-8C
Gly-7C
Gly-9C
(B)
-0.61
-0.61
-0.68
-0.77
-0.77
-0.79
Terminal
-0.3
+0.3
Electrostatic Potential
(C)
Terminal
HOMO
LUMO
131.77 Kcal/mol
59.86 Kcal/mol
(D)
Gly-98H
Ser-99H
Gly-96H
Gly-96L
Thr-97H
Terminal
-0.3
+0.3
Electrostatic Potential

Fig. S22.

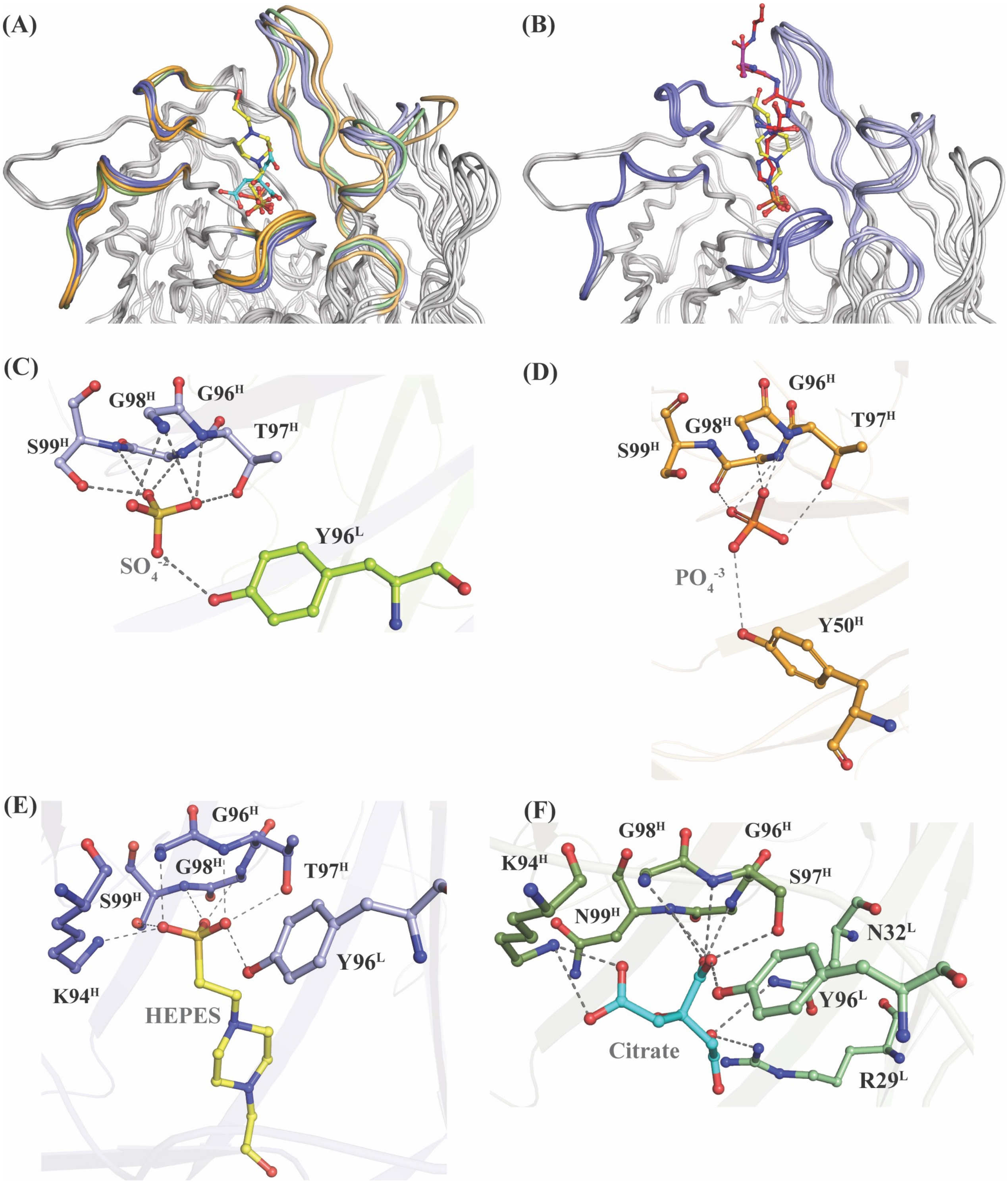

(A)
(B)
(C)
G98H
G96H
S99H
T97H
Y96L
SO4-2
(D)
G96H
G98H
T97H
S99H
PO4-3
Y50H
(E)
G96H
G98H
T97H
S99H
K94H
Y96L
HEPES
(F)
G98H
G96H
K94H
S97H
N99H
N32L
Y96L
Citrate
R29L

**Fig. S23.**

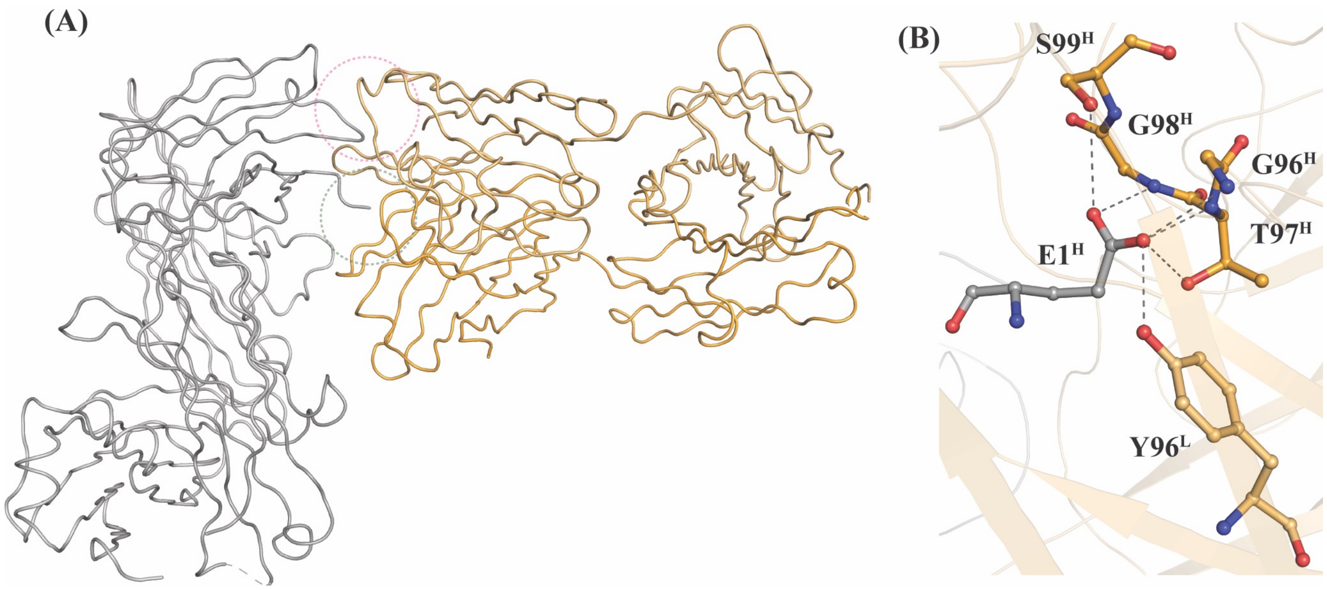

### Table S1.

**Table S1.** Mutagenic oligonucleotides used to generate libraries hSC44 1-6.

| Name | Mutagenic Oligonucleotide Sequence |
|---|---|
| hSC44.L1.1 | CGTGCCAGTCAGTCCGTG(N4:10101070)(N3:10107010)(N3)(N2:10701010)(N3)(N4)AAC(N1:70101010)(N1)(N1)(N1)(N1)(N2)GTAGCCTGGTATCAACAG |
| hSC44.L1.2 | CGTGCCAGTCAGTCCGTG(N4:10101070)(N3:10107010)(N3)CGTAACAAG(N1:70101010)(N1)(N2:10701010)GTAGCCTGGTATCAACAG |
| hSC44.L3.1 | TACTGTGTGGGC(N2:10701010)(N1:70101010)(N4:10101070)TATGGCAGCGAAAACGATGCGTAT(N4)(N1)(N2)GCGTTCGGACAGGGTACC |
| hSC44.L3.2 | TACTGTGTGGGC(N2:10701010)(N1:70101010)(N4:10101070)TAT(N3:10107010)(N3)(N2)(N1)(N3)(N2)(N3)(N1)(N1)(N1)(N1)(N2)(N3)(N1)(N4)(N3)(N2)(N3)TAT(N4)(N1)(N2)GCGTTCGGACAGGGTACC |
| hSC44.H1.1 | GGCTTCAGCATTGATAGC(N4:10101070)(N1:70101010)(N4)(N3:10107010)(N3)(N2:10701010)TTTAGCTGGGTGCGTCAGGCC |
| hSC44.H2.1 | CTGGAACATATTGGC(N4:10101070)(N1:70101010)(N4)CTG(N1)(N2:10701010)(N2)(N3:10107010)(N2)(N2)GGCGGCCGTGCGTTTTATGCC |
| hSC44.H2.2 | CTGGAACATATTGGCTATCTG(N1:70101010)(N2:10701010)(N2)GCGGGCGGC(N2)(N3:10107010)(N4:10101070)GCGTTTTATGCCAGCTGG |
| hSC44.H3.1 | GTCTATTATTGTGCT(N1:70101010)(N1)(N1)(N2:10701010)(N4:10101070)(N4)(N3:10107010)(N3)(N2)(N1)(N3)(N2)(N3)(N3)(N2)(N1)(N1)(N2)(N2)(N2)(N3)(N3)(N4)(N4)GCGATTTGGGGTCAAGGAACC |
| hSC44.H3.2 | GTCTATTATTGTGCT(N1:70101010)(N1)(N3:10107010)(N2:10701010)(N4:10101070)(N4)(N3)(N3)(N2)(N1)(N3)(N2)GGC(N1)(N1)(N2)CCG(N3)(N4)(N4)GCGATTTGGGGTCAAGGAACC |
| hSC44.V.1 | AGCATTGATAGCTATGGC(N4:10101070)(N4)(N4)(N1:70101010)(N3:10107010)(N2:10701010)TGGGTGCGTCAGGCCCCG |
| hSC44.V.2 | GGTAAGGGCCTGGAA(N2:10701010)(N1:70101010)(N4:10101070)(N1)(N4)(N4)GGC(N4)(N1)(N4)CTGACCGCGGGC |
| hSC44.V.3 | GGCCGTGCG(N4:10101070)(N4)(N4)TATGCC(N1:70101010)(N3:10107010)(N2:10701010)(N4)(N3)(N3)(N3)(N2)(N2)AAG(N1)(N3)(N2)CGT(N1)(N3)(N2)ACTATAACC(N2)(N3)(N4)AACACAAAC |
| hSC44.V.4 | GAAAACACA(N3:10107010)(N4:10101070)(N4)ACCCTAAAAATGAACAGC |

**Table S2.**

**Table S2.** Final concentrations of reagents used on various days of affinity selections.

| | Day 1 | Day 2 | Day 3 | Day 4 | Day 5 |
|---|---|---|---|---|---|
| BSA† | 1% | 1% | 1% | 1% | 1% |
| Streptavidin† | 10µg/mL | - | 10µg/mL | - | 10µg/mL |
| Neutravidin† | - | 10µg/mL | - | 10µg/mL | - |
| No $PO_4^-$ Peptide*,† | - | 10nM | 25nM | 50nM | 100nM |
| 1-pTza Peptide*,† | - | 1nM | 5nM | 10nM | 20nM |
| 3-pTza/pHis Peptide*,‡ | 100nM | 100nM | 50nM | 25nM | 10nM |
| No. Wells | 24 | 12 | 8 | 6 | 4 |
| No. Washes | 4 | 6 | 8 | 10 | 12 |

†Negative selections, ‡Positive Selections

*Captured using 2µg/mL streptavidin/neutravidin on alternating days

### Table S3.

**Table S3**. Elbow angles of variants of hSC44 Fabs

| Fab | Ligand | Protomers | Elbow angle (°) |
|---|---|---|---|
| hSC44.S1C | ACLYana-3-pTza peptide | HL | 138.9 |
| | | AB | 157.3 |
| hSC44.S1C | ACLYana-3-pHis peptide | HL | 137.8 |
| | | AB | 157.1 |
| hSC44.S1C | - | HL | 138.9 |
| | | AB | 159.7 |
| hSC44.S1C.20 | ACLYana-3-pTza peptide | HL | 135.9 |
| hSC44.S1C.20 | ACLYana-3-pHis peptide | HL | 135.8 |
| hSC44.S1CE.20 | - | HL | 165.3 |
| hSC44.S1CE.20 | - | HL | 172.6 |
| hSC44.S1C.20.N32F$^{L}$ | ACLYana-3-pTza peptide | HL | 165.1 |
| | | AB | 147.9 |
| hSC44.S1C.20.N32F$^{L}$ | ACLYana-3-pHis peptide | HL | 165.4 |
| | | AB | 149.0 |
| hSC44.S1C.20.N32F$^{L}$ | - | HL | 156.2 |
| hSC44.S1CE.20.N32F$^{L}$ | - | HL | 156.1 |

### Table S4.

#### Table S4. Data collection and refinement statistics for newly solved Fab-Antigen crystal structures.

**Table S4.** X-ray data collection and refinement statistics for newly solved Fab-Antigen structures.

| **Structure Name** | **hSC44.S1C** | **hSC44.S1C** | **hSC44.S1C** | **hSC44.S1C.20** | **hSC44.S1C.20** | **hSC44.S1CE.20** | **hSC44.S1CE.20** | **hSC44.S1C.20.N32F[L]** | **hSC44.S1C.20. N32F[L]** | **hSC44.S1C.20.N32F[L]** | **hSC44.S1CE.20.N32F[L]** |
|---|---|---|---|---|---|---|---|---|---|---|---|
| **Combining site ligand** | **AGAG-3pTza-AGAG** | **AGAG-3pHis-AGAG** | **No peptide (citrate)** | **AGAG-3pTza-AGAG** | **AGAG-3pHis-AGAG** | **No peptide (GluH1')** | **No peptide (PO4)** | **AGAG-3pTza-AGAG** | **AGAG-3pHis-AGAG** | **No peptide (SO4)** | **No peptide (HEPES)** |
| **Data collection** | | | | | | | | | | | |
| Beamline | SSRL 12-1 | SSRL 12-1 | APS 23-ID-B | APS 23-ID-D | SSRL 12-1 | ALS 5.0.1 | ALS 5.0.1 | ALS 5.0.1 | ALS 5.0.1 | ALS 5.0.1 | ALS 5.0.1 |
| Wavelength (Å) | 0.97946 | 0.97946 | 1.03317 | 1.0332 | 0.97946 | 0.97741 | 0.97741 | 0.97741 | 0.97741 | 0.97741 | 0.97741 |
| Resolution (Å)[a] | 39.34-1.95 | 38.72-2.20 | 46.91-1.75 | 41.04-1.85 | 34.82-1.90 | 47.17-2.45 | 49.79-2.40 | 47.25-1.98 | 47.19-2.09 | 48.05-1.94 | 46.27-1.84 |
| Space group | $P2_12_12$ | $P2_12_12$ | $P2_12_12$ | C2 | $P2_1$ | $P2_12_12_1$ | $P4_32_12$ | $P2_12_12_1$ | $P2_12_12_1$ | $P2_12_12_1$ | $P2_12_12_1$ |
| Unit cell (Å) | 92.50, 139.62, 73.62 | 93.25, 139.06, 73.71 | 93.82, 138.85, 73.21 | 79.96, 73.12, 88.07 | 44.11, 72.61, 70.33 | 49.7, 73.3, 148.5 | 72.3, 72.3, 205.9 | 71.55, 73.95, 239.65 | 71.28, 74.03, 239.28 | 69.47, 73.25, 96.11 | 48.57, 72.07, 152.2 |
| ( °) | 90, 90, 90 | 90, 90, 90 | 90, 90, 90 | 90, 111.24, 90 | 90, 98.21, 90 | 90, 90, 90 | 90, 90, 90 | 90, 90, 90 | 90, 90, 90 | 90, 90, 90 | 90, 90, 90 |
| Total reflections | 733,426 | 637,037 | 688,286 | 155,586 | 148,666 | 127,101 | 219,439 | 538,821 | 418,880 | 284,579 | 293,288 |
| Unique reflections | 68,051(3151) | 46,924(2432) | 95,210(4089) | 40,163(1984) | 33,299(1333) | 19,822(839) | 22,383(1088) | 89,758(4417) | 71,747(3417) | 36,687(1806) | 46,607(2191) |
| Multiplicity | 10.8(7.8) | 12.9(9.4) | 7.2(5.5) | 3.9(3.3) | 4.5(3.6) | 6.4(5.7) | 9.8(6.2) | 6.0(5.6) | 5.8(5.2) | 7.8(7.6) | 6.3(4.5) |
| Completeness (%) | 97.5(91.3) | 99.9(99.9) | 98.1(85.3) | 99.8(99.4) | 96.1(77.1) | 96.3(84.2) | 100(99.7) | 100(100) | 95.2(92.7) | 100(100) | 99.2(94.4) |
| Mean (I)/ $\sigma_I$ | 13.1(1.7) | 11.8(1.8) | 10.0(1.6) | 6.9(1.1) | 9.9(2.8) | 19.1(2.1) | 28.0(2.7) | 8.9(1.5) | 13.9(1.5) | 11.4(1.0) | 16.0(1.2) |
| $R_{merge}$[b](%) | 14.3(193) | 22.1(483) | 12.4(117) | 13.1(128) | 16.2(120) | 6.3(85.4) | 7.1(76.1) | 12.5(135) | 10.1(130) | 18.0(238) | 10.0(178) |
| $R_{meas}$[c](%) | 14.9(205) | 23.0(512) | 13.3(129) | 15.1(151) | 18.3(141) | 6.9(94.1) | 7.5(83.0) | 13.7(150) | 11.0(144) | 19.3(254) | 10.8(200) |
| $R_{pim}$[d](%) | 4.4(67.3) | 6.4(166) | 4.9(52.9) | 7.4(79.2) | 8.2(71.9) | 2.7(38.7) | 2.3(32.2) | 5.6(63.6) | 4.4(61.1) | 6.7(88.4) | 4.2(90.4) |
| $CC_{1/2}$[e](%) | 99.8(47.2) | 99.8(56.5) | 99.7(41.1) | 99.5(44.1) | 99.2(55.0) | 99.9(70.6) | 99.9(80.2) | 99.5(39.6) | 99.7(43.2) | 99.3(37.4) | 99.6(37.1) |
| **Refinement** | | | | | | | | | | | |
| Refinement resolution (Å)[a] | 39.34-1.95 | 38.72-2.20 | 46.91-1.75 | 41.04-1.85 | 34.82-1.90 | 47.17-2.45 | 49.79-2.40 | 47.25-1.98 | 47.2-2.09 | 48.05-1.94 | 46.27-1.84 |
| # reflections in refinement (work/free) | 64,623/3393 | 46,924/2489 | 90,339/4802 | 38,147/1978 | 31,562/1721 | 18,793/985 | 21,221/1079 | 85,368/4291 | 68,152/3509 | 34,775/1846 | 44,224/2299 |
| $R_{work}/R_{free}$ (%) | 21.2/26.0 | 23.0/27.5 | 21.0/23.9 | 20.5/24.3 | 17.2/21.3 | 24.4/29.0 | 25.8/29.3 | 22.4/25.7 | 21.8/24.5 | 21.8/24.7 | 20.4/23.6 |
| # atoms (Fab/Peptide/Solvent) | 6556/64/223 | 6556/69/64 | 6532/na/476 | 6514/57/283 | 6526/69/188 | 3286/na/10 | 3287/na/34 | 6589/64/206 | 6552/72/334 | 3278/na/149 | 3336/na/480 |
| RMS (bonds) | 0.006 | 0.002 | 0.006 | 0.003 | 0.005 | 0.005 | 0.003 | 0.008 | 0.002 | 0.003 | 0.005 |
| RMS (angles) | 0.91 | 0.6 | 0.83 | 0.75 | 0.77 | 0.71 | 0.61 | 0.94 | 0.52 | 0.66 | 0.86 |
| Ramachandran favoured/allowed/ outliers (%) | 96.9/2.9/0.2 | 97.6/2.4/0 | 96.7/2.9/0.4 | 97.7/2.3/0 | 97.7/2.1/0.2 | 95.1/4.7/0.2 | 96.0/4.0/0 | 96.4/3.6/0 | 97.3/2.6/0.1 | 97.7/2.3/0 | 98.4/1.6/0 |
| Ramachandran plot Z score | 0.02 | -0.1 | 0.2 | -0.1 | 0.8 | -1.4 | -1.6 | -0.29 | -0.81 | 0.06 | 0.39 |
| Clashscore[f] | 3.1 | 1.8 | 3.7 | 0.6 | 1.5 | 3.2 | 4.6 | 2.1 | 1.4 | 2.6 | 3.5 |
| Wilson B ($Å^2$) | 30 | 41 | 22 | 22 | 21 | 53 | 51 | 32 | 36 | 27 | 25 |
| Average B ($Å^2$) for all atoms/Fab/ Peptide/Solvent | 40/38/54/34 | 56/56/85/43 | 34/34/na/30 | 29/29/38/32 | 25/25/22/27 | 63/63/na/46 | 72/72/na/51 | 42/42/56/36 | 43/43/56/41 | 30/30/46/29 | 31/30/20/36 |
| PDB ID | **8UJI** | **8UIT** | **8UIO** | **8UIH** | **8UIG** | **8UHT** | **8UHS** | **8UHP** | **8UHN** | **8UHJ** | **8UHH** |

[a]Numbers in parentheses are for highest resolution shell

[b]$R_{merge} = \Sigma_{hkl} \Sigma_{i=1,n}|I_i(hkl) - \langle I(hkl)\rangle| / \Sigma_{hkl} \Sigma_{i=1,n} I_i(hkl)$

[c]$R_{meas} = \Sigma_{hkl} \sqrt{(n/n-1)} \Sigma_{i=1,n}|I_i(hkl) - \langle I(hkl)\rangle| / \Sigma_{hkl} \Sigma_{i=1,n} I_i(hkl)$

[d]$R_{pim} = \Sigma_{hkl} \sqrt{(1/n-1)} \Sigma_{i=1,n}|I_i(hkl) - \langle I(hkl)\rangle| / \Sigma_{hkl} \Sigma_{i=1,n} I_i(hkl)$

[e]$CC_{1/2}$ = Pearson Correlation Coefficient between two random half datasets

[f]Number of unfavorable all-atom steric overlaps ≥ 0.4Å per 1000 atoms

#### Table S5.

**Table S5.** Constructs with ligand, crystallization conditions and cryoprotectant used for crystallization experiments

| pHis Fab | Ligand | Crystallization condition | Cryoprotectant |
|---|---|---|---|
| hSC44.S1C (C1) | AGAG-3-pTza-AGAG | 0.2 M Lithium citrate, 20% PEG3350 | 30 % Ethylene glycol |
| hSC44.S1C (A7) | AGAG-3-pHis-AGAG | 0.2 M Calcium chloride, 20% PEG3350 | 30 % Ethylene glycol |
| hSC44.S1C (A2) | No ligand | 0.2 M Lithium citrate, 20% PEG3350 | 30 % Ethylene glycol |
| hSC44.S1C.20 (J13) | AGAG-3-pTza-AGAG | 0.2 M tri-potassium citrate, 20% PEG3350 | 25% PEG400 |
| hSC44.S1C.20 (B5) | AGAG-3-pHis-AGAG | 0.2 M tri-potassium citrate, 20% PEG3350 | 25% PEG400 |
| hSC44.S1CE.20 (C16) | No ligand | 0.1 M Tris pH 8.5, 8% PEG8000 | 30 % Ethylene glycol |
| hSC44.S1CE.20 (A10) | No ligand | 0.1 M HEPES pH 7.5, 20% PEG4000, 10% 2-propanol | 30 % Ethylene glycol |
| hSC44.S1C.20.N32F$^{L}$ (E12) | AGAG-3-pTza-AGAG | 0.08 M Sodium Cacodylate pH 6.5, 0.16 M Calcium acetate, 20% glycerol, 14.4% PEG8000 | 10% glycerol |
| hSC44.S1C.20.N32F$^{L}$ (E14) | AGAG-3-pHis-AGAG | 0.08 M Sodium Cacodylate pH 6.5, 0.16 M Calcium acetate, 20% glycerol, 14.4% PEG8000 | 10% glycerol |
| hSC44.S1C.20.N32F$^{L}$ (C8) | No ligand | 0.1 M Tris pH 8.5, 0.2 M Lithium sulfate, 40% PEG400 | Well solution |
| hSC44.S1CE.20.N32F$^{L}$ (C15) | No ligand | 0.1 M HEPES pH 7.5, 20% PEG4000, 10% 2-propanol | 30 % Ethylene glycol |

### Table S6.

**Table S6**: Changes in the ligand binding free energies caused by the N32F$^L$ mutation. The first column shows the binding free energy differences from FEP calculations ($\Delta\Delta \boldsymbol{G}_{\boldsymbol{bind}}^{\boldsymbol{WT}\rightarrow\boldsymbol{M}}$) of 3-pHis, whereas the last column shows the experimentally obtained binding free energy differences.

| System | Relative Binding Free energy $\Delta\Delta G_{bind}$ (Kcal/mol) | $K_D$(M) | Experimental binding free energy $\Delta\Delta G_{bind}$ (Kcal/mol) |
|---|---|---|---|
| pHis_SC44H.20.F (N32F$^L$) | -1.1±0.4 | 1.68E-09 | $RT \ln \frac{K_D^M}{K_D^{WT}}$ =-1.4 |
| pHis_SC44H.20.F (WT) | | 2.10E-08 | |
